# KnowVolution of an efficient polyamidase through molecular dynamics simulations of incrementally docked oligomeric substrates

**DOI:** 10.1101/2024.08.13.607760

**Authors:** Hendrik Puetz, Alexander-Maurice Illig, Mariia Vorobii, Christoph Janknecht, Francisca Contreras, Fabian Flemig, Ulrich Schwaneberg

## Abstract

Management of synthetic polymer waste is one of the most pressing challenges for society today. Enzymatic recycling of polycondensates like polyamides (PA), however, remains limited due to a lack of efficient polyamidases. This study reports the directed evolution of the polyamidase NylCp2-TS. Key positions involved in enzyme-substrate interactions and PA 6 hydrolysis are identified through random mutagenesis and molecular dynamics (MD) simulations. The final variant, NylC-HP (NylCp2-TS^F134W/D304M/R330A^), exhibits a 6.9-fold increased specific activity (520 ± 19 µmol6-AHAeq. h^−1^ mgenzyme^−1^) and enhanced thermal stability (*T*m = 90 °C, Δ*T*m = 4.2 °C), making NylC-HP the fastest polyamidase for PA 6 and PA 6,6 hydrolysis. Despite the improved reaction rate, the degree of depolymerization remains below 1 %. To understand the molecular basis of achieved improvements and factors limiting the degree of depolymerization, intra- and intermolecular interactions of various enzyme-substrate complexes are analyzed by incremental docking of PA 6 tetramers and MD simulations. After optimizing the activity and stability of NylC-HP, the findings suggest that widening the substrate binding pocket is likely necessary to improve substrate accessibility to target more buried attack sites on the polymer surface, and thereby enhancing the degree of depolymerization.

## 1 Introduction

Enzymatic polymer recycling holds the potential to play a pivotal role in the establishment of a circular plastic economy. Consequently, efforts to industrially scale the depolymerization of PET (polyethylene terephthalate) have been actively pursued in recent years. PETases could be applied to completely depolymerize PET at industrially relevant solids loading bringing cost-effective PET recycling into reach.^1-3^ This was greatly fueled by enzyme engineering campaigns augmenting the stability, specific activity, and the degree of depolymerization of PETases.^2, 4, 5^ Recently, there has been a growing interest from academia and industry in enzymatic recycling of PA (polyamide) and PUR (polyurethane).^6–11^ To this date, PA recycling processes are limited by the availability of depolymerases with sufficient performance. To realize a circular polyamide economy, sustained efforts in protein engineering of known depolymerases and the discovery of novel biocatalysts are indispensable.^11–14^

As of now, the nylon hydrolase NylCp2 and its isoforms stand out as the only hydrolases to considerably depolymerize PA 6 and PA 6,6. NylCp2 has already undergone rational engineering that resulted in the remarkably stable polyamidase NylCp2-TS.^15, 16^ Despite successfully enhancing the melting temperature (*T*m) to 88 °C, this achievement was accompanied by significant decrease in the turnover frequency.^16^ Consequently, the next step to advance enzymatic PA recycling is to design a polyamidase with activity and degree of depolymerization comparable to PETases used at industrial-scale PET recycling (963 µmolTPAeq. h^−1^ mg ^−1^ ; 98 % in 24 h).^1^ In a recent study, we developed and validated a high-throughput amine screening system for directed polyamidase evolution (AMIDE).^12^ In this study, we applied AMIDE, combined with incremental substrate docking and molecular dynamics (MD) simulations, to engineer NylCp2-TS in a KnowVolution campaign.^17^

KnowVolution is a versatile protein engineering strategy that has been used extensively to improve enzyme traits such as activity; thermal, ionic liquid and pH resistance; substrate affinity; and substrate specificity.^17–22^ KnowVolution combines computational analyses with directed evolution to optimize properties with minimized experimental efforts and maximized improvements. The KnowVolution strategy comprises four phases: (I) identification of beneficial amino acid positions by screening random mutagenesis or smart libraries, (II) determination of beneficial substitutions for previously identified positions, (III) selection of positions to recombine concerning their epistasis, and (IV) recombination of beneficial substitutions to maximize enzyme fitness.

Substrate docking is important for determining and evaluating the pose of a ligand (e.g., substrate) in complex with a receptor (e.g., enzyme) for applications such as virtual drug screening.^23^ Since docking is optimized for predicting the bound conformation and free energy of small molecules to the receptor, incremental approaches are necessary to extend its applicability to large molecules with many rotatable bonds such as peptides.^24^ Recently, docking and subsequent molecular dynamics simulations of a small polymer model substrate (2PET) and a polymer-degrading enzyme (PETase) were applied to decipher the effects of substitutions on enzyme activity at the molecular level.^25^ Aliphatic polycondensates (e.g., PA 6, PA 6,6, PLA) display a significantly higher number of rotatable bonds than PET. To enable the investigation of the interactions between NylCp2-TS and PA 6, we implemented an incremental docking workflow using oligomeric PA 6 substrates. Furthermore, MD simulations were performed to model the dynamic behavior of ligand-receptor complexes.^26^

In this study, we present the first directed evolution campaign for a polyamidase and employed the KnowVolution strategy. We performed random mutagenesis and MD simulations of the enzyme-substrate complex to identify key positions crucial for the interaction of NylCp2-TS with PA 6 and PA 6,6, and its hydrolysis. Recombination of the identified beneficial substitutions enabled us to enhance the specific activity of NylCp2-TS by up to 6.9-fold. The finally obtained NylC-HP variant is the highest-performing polyamidase to date in respect to PA 6 and PA 6,6 depolymerization efficiency. Our study highlights the potential of enzyme engineering to enhance enzymatic polyamide hydrolysis while also identifying limitations that need to be addressed in future efforts. Specifically, for NylC-HP, we propose that significantly widening the substrate binding pocket is necessary to achieve higher degrees of depolymerization.

## 2 Results and discussion

The goal of this study is to generate knowledge to enable efficient engineering of polyamidases for PA 6 and PA 6,6 hydrolysis. We focused on enhancing the PA 6 depolymerization efficiency of NylCp2-TS and understanding the enzyme-polymer interactions within the binding pocket that govern substrate hydrolysis. These aspects are described in three sections. First, KnowVolution is applied to engineer NylCp2-TS towards enhanced specific activity. Next, we experimentally characterize the highest-performing variant (NylC-HP). Finally, we conduct computational analyses to identify changes in enzyme-enzyme and enzyme-substrate interactions that lead to improved substrate hydrolysis and provide molecular understanding on structural limitations for PA degradation.

Enzymatic PA recycling is an important research area for establishing a circular economy for synthetic polyamides and is gaining attention from academic research groups and industry.^9–11, 27^ Using characterized polymers as substrates and applying defined pretreatment techniques is essential for ensuring comparability between studies conducted by different research groups, as demonstrated in enzymatic PET degradation research.^1^ We observed background product formation in PA 6 and PA 6,6 degradation studies, primarily due to the extraction of unpolymerized monomers and the spontaneous hydrolysis of 6-aminohexanoic acid (6-AHA) in the case of PA 6. In this study, all results were background-corrected to monitor enzyme net activity. We aimed to maximize the reproducibility and comparability of this study. Thus, we utilized untreated Gf-PA 6 film (Goodfellow GmbH, 0.2 mm thickness; *X*c,PA 6 = 21 %; *M*w,PA 6 = 108.5 kDa; *M*n,PA 6 = 44.52 kDa) and Gf-PA 6,6 granules (Goodfellow GmbH, 3 mm diameter; *X*c,PA 6,6 = 39 %; *M*w,PA 6,6 = 61.84 kDa; *M*n,PA 6,6 = 29.30 kDa) and characterized them in respect to their crystallinity and molecular weight (**Figure S1 and Table S1-S2**). Recrystallization of PET during the enzymatic hydrolysis process significantly reduces depolymerization efficiency.^1, 4^ Thus, we also determined the extent of PA 6 and PA 6,6 recrystallization during heat treatment mimicking enzymatic depolymerization process conditions. In a previous study, slight increases in crystallinity of PA 6 were observed when incubated at 60 °C and 70 °C for up to 10 days.^10^ From an economic perspective, it is unlikely to extend the process to several days.^28, 29^ To determine maximum process temperatures, we analyzed the crystallinity of Gf-PA 6 film and Gf-PA 6,6 granules after incubation at temperatures ranging from 40 °C to 90 °C for 24 h and confirmed that no significant increases in crystallinity occurred, even at 90 °C (**Figure S1A,B and Table S1**). These findings suggest that, unlike enzymatic PET hydrolysis, both PA 6 and PA 6,6 depolymerization are not limited by recrystallization above the glass transition temperature (*T*g, PA 6 = 53 °C, *T*g, PA 6,6 = 57 °C).^30^ Therefore, PA depolymerization can be conducted at temperatures up to at least 90 °C, provided that appropriately thermostable enzymes are available.

### 2.1 MD simulation-guided directed evolution of NylCp2-TS

We sought to increase the depolymerization efficiency of NylCp2-TS towards PA 6 by combining random mutagenesis and rational design (i.e., active site engineering performing docking and MD simulations) as part of a four-step KnowVolution campaign.^17^ The specific activity of depolymerases is an established performance parameter and was used for evaluating and selecting NylCp2-TS variants.^2, 4, 31^

#### 2.1.1 Identification of potentially beneficial positions through random mutagenesis and MD simulations

The first step of the KnowVolution strategy comprises the identification of potentially beneficial positions. We applied random mutagenesis to identify beneficial positions across the entire gene and MD simulations of the enzyme-substrate complex to determine positions involved in enzyme-substrate interactions. From screening a random mutagenesis library comprising 1700 clones, we identified eleven potentially beneficial positions (V18, L24, P27, F38, D99, F100, G111, A160, D209, F301, and D304). For *in silico* identification, we determined the contact frequency of the individual residues of the enzyme with the substrate observed in the MD simulations. To generate the enzyme-substrate complexes, the PA 6 tetramers Ace-[6-AHA]4-COO^−^ and NMe-[6-AHA]4-NH ^+^; comprising four repeating units of 6-AHA, one capped end representing the continuation of the polymer chain, and one charged end representing the terminus of the polymer chain; were incrementally docked into the binding pocket of NylCp2-TS (**Figure 1 and Figure 2A**). We observed two substrate orientations with the tetramer’s attack site near the active site, but with mirrored substrate termini (**Figure S2**).

**Figure 1.**
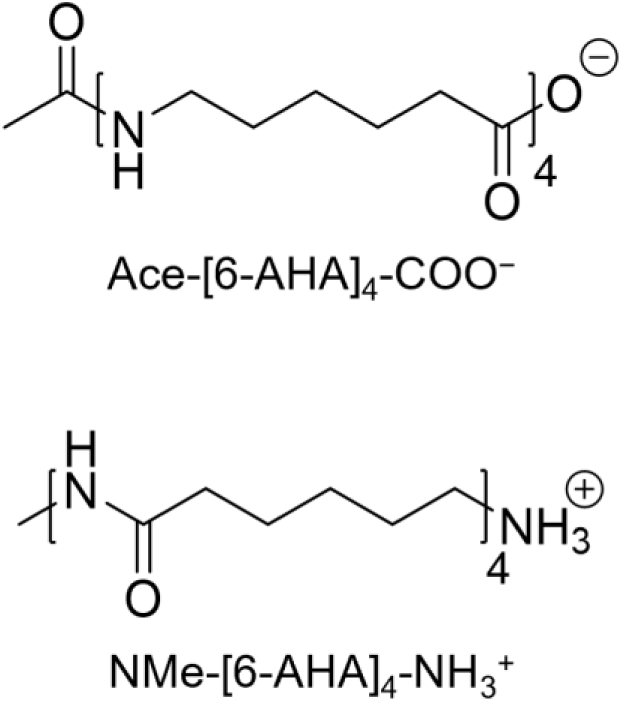
PA 6 tetramers used for substrate docking and MD simulation. Ace-[6-AHA]_4_-COO^−^ was used as proxy for the PA 6 polymer chain carboxy-terminus and NMe-[6-AHA]4-NH3^+^ for the amine-terminus. The opposing termini were capped to mimic the continuation of the polymer chain.

**Figure 2.**
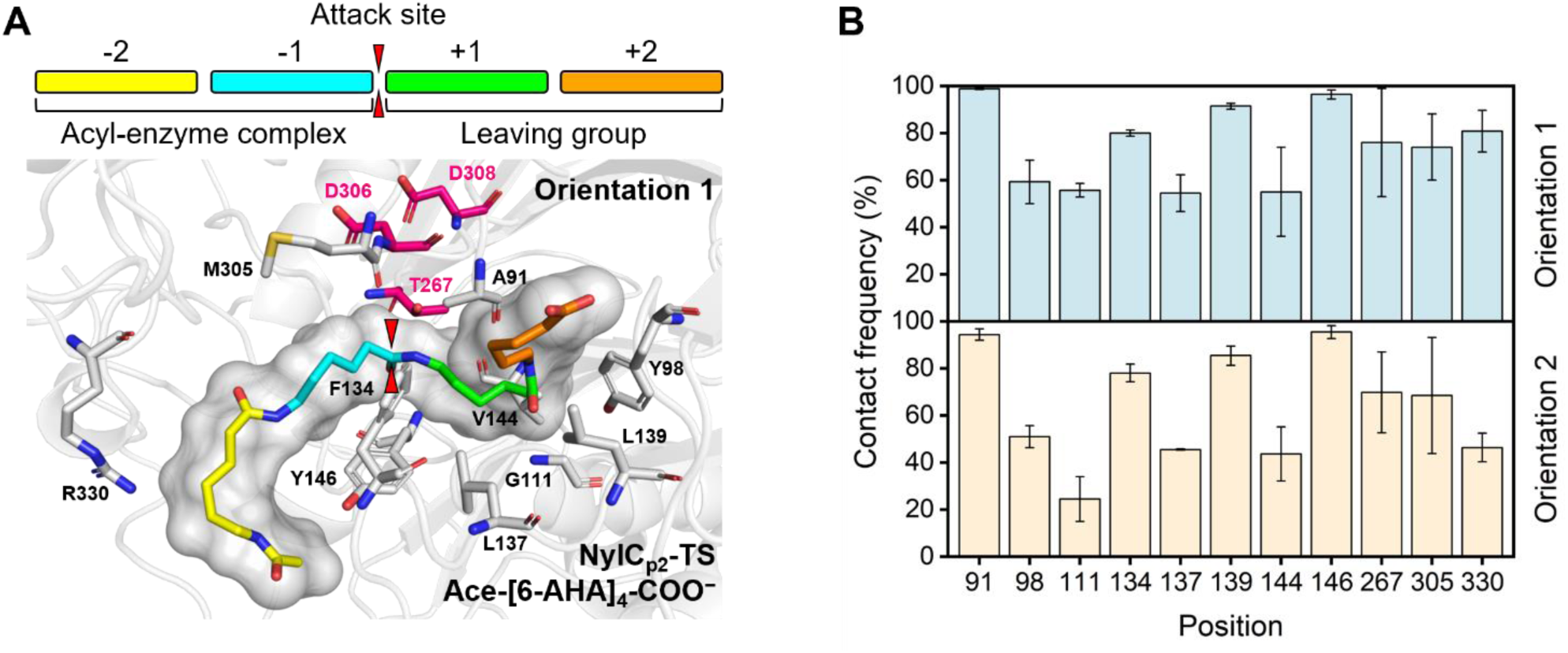
Identification of the substrate interaction sphere of NylCp2-TS and PA 6. A) Representative structure of enzyme-substrate complex NylCp2-TS/Ace-[6-AHA]_4_-COO^−^(orientation 1). Residues with ≥ 50 % of the simulation time in ≤ 2.5 Å contact sphere of the enzyme are shown as sticks. The catalytic triad is highlighted in pink. B) Contact frequencies of the ≤ 2.5 Å contact sphere of NylCp2-TS and both Ace-[6-AHA]_4_-COO^−^ orientations.

NylCp2-TS exhibited a lower free energy of binding for Ace-[6-AHA]4-COO^−^ in both orientations (orientation 1: Ace-[6-AHA]4-COO^−^ and orientation 2: ^−^OOC-[6-AHA]4-Ace) compared to NMe-[6-AHA]4-NH3^+^ (**Figure S2 and Table S3**). Therefore, MD simulations of the enzyme-substrate systems NylCp2-TS/Ace-[6-AHA]4-COO^−^ (orientation 1) and NylCp2-TS/^−^OOC-[6-AHA]4-Ace (orientation 2) were performed. We considered a residue to be in contact with the substrate when the distance between any atom of the two was less than or equal to 2.5 Å. **Figure 2A** displays a representative structure of the NylCp2-TS/Ace-[6-AHA]4-COO^−^ enzyme-substrate complex, with interacting residues highlighted. The reported contact frequencies represent the mean of three independent 150 ns simulation runs and give the percentage of simulation time of existing contacts (**Figure 2B**). Positions that were in contact with the substrate for at least 50 % of the simulation time were selected for experimental testing. The MD simulations revealed ten positions (A91, Y98, G111, F134, L137, L139, V144, Y146, T267, M305, and R330) to be in the 2.5 Å interaction sphere for at least 50 % of the simulation time (**Figure 2A,B**). Even though T267 showed an average contact frequency of ≥ 50 %, it was not considered for site-saturation, as it acts as nucleophilic residue in the catalytic triad and is deemed indispensable for substrate hydrolysis.^16, 32^

#### 2.1.2 Determining beneficial substitutions through site-saturation mutagenesis

In the second step, site-saturation mutagenesis libraries for 20 positions (i.e., V18, L24, P27, F38, A91, Y98, D99, F100, G111, F134, L137, L139, V144, Y146, A160, D209, F301, D304,

M305, and R330) were generated using degenerate primers (NNK) and screened with Gf-PA 6 film as substrate via the AMIDE method to identify substitutions leading to improved specific activity.^12^ Specifically, we quantified 6-AHA equivalents (primarily 6-AHA monomers and dimers) by labelling released amines, serving as indicators of hydrolysis reactions catalyzed by the enzyme. Out of 20 positions, substitutions at five positions (D99, F134, F301, D304, and R330) showed significantly improved activity (1.5 to 3.0-fold) compared to NylCp2-TS (**Figure 3**). For D99(R/G/V), F134W, F301L, and R330(A/Q) a small set of 1-3 amino acid substitutions showed improved activity. Interestingly, position 304 exhibited significant variability, as many substitutions D304(M/E/Q/L/V/R/W) resulted in enhanced PA 6 hydrolysis activity.

**Figure 3.**
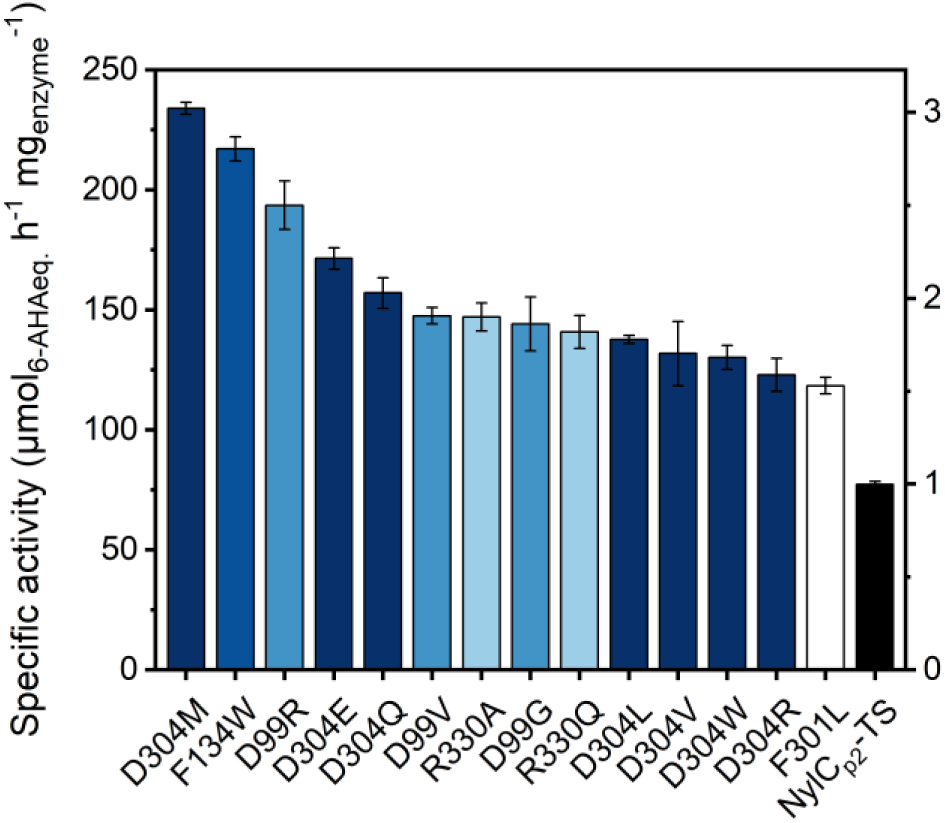
Specific activities of NylCp2-TS single substitutions. Specific activities were calculated from released 6-AHA equivalents after incubation with Gf-PA 6 film (50 nM/1.90 mg L^−1^ enzyme, 456 g L^−1^ Gf-PA 6, 50 mM bicine, pH 8.0, 100 mM NaCl, 60 °C, 2 h). Reactions were conducted in at least duplicate. Error bars represent the standard error of the mean.

#### 2.1.3. Selection of positions for recombination

In the third step, we selected positions for which we determined beneficial substitutions for recombination. The recombination of beneficial substitutions can lead to negative epistasis, resulting in only moderate improvements or even a reduction in fitness, when individually advantageous substitutions do not exhibit additive effects.^33^ Epistasis of recombined amino acid substitutions is more likely to occur when positions are in close spatial proximity, as in the case of F301 and D304.^34^ To assess epistasis, we evaluated the compatibility of substitutions at these positions. We screened an iterative site-saturation mutagenesis library where D304 was saturated using NylCp2-TS^F301L^ as the parental enzyme. This resulted in negative epistasis for all F301L/D304X double-substituted variants. Due to negative epistasis and the fact that F301L displayed only a weak beneficial effect on the activity of NylCp2-TS (1.5-fold ± 0.1 improved specific activity), we excluded F301L and continued recombination with beneficial substitutions at the remaining positions (i.e., D99, F134, D304, and R330).

#### 2.1.4 Recombination of determined beneficial substitutions yields a polyamidase with high performance

In the fourth and final step, we recombined beneficial positions to maximize the activity of NylCp2-TS. The number of beneficial substitutions at each position differed greatly, and most prominently, seven substitutions at position D304 increased the specific activity of NylCp2-TS significantly (1.6 to 3.0-fold) (**Figure 3**). The variant with the substitution F134W exhibited the second-highest catalytic improvement, with a single beneficial phenylalanine-to-tryptophan substitution increasing the specific activity (2.8-fold ± 0.1). Substitutions at D99(R/G/V) and R330(A/Q) enhanced the specific activity, with three substitutions at position D99 (1.9 to 2.5- fold) and two at position R330 (1.8 to 1.9-fold). The best single substitution D304M resulted in a 3.0-fold ± 0.0 improved activity. Introducing methionine at position D304 may, however, not necessarily show the highest synergy when recombined with F134W. Hence, iterative site saturation is advisable wherein variants with improved activity are used as a new reference point for another round of screening.^35, 36^ Accordingly, we site-saturated D304 with NylCp2-TS^F134W^ as the parental enzyme to trace the NylCp2-TS^F134W/D304X^ double-substituted variant with the highest specific activity. 45 clones exhibiting higher activity than the parental enzyme were sequenced. Among them, NylCp2-TS^F134W/D304M^ was the most frequently occurring variant (13%) and showed the most improved specific activity for PA 6 (5.3-fold ± 0.1) (**Figure 4, Figure S3, and Table S4-S5**). Following this double substitution, eleven recombined variants with D99(R/G/V) and R330(A/Q) remained for evaluation. We generated all possible triple and quadruple substitutions based on NylCp2-TS^F134W/D304M^ (V1-V11, **Table S5**) and determined respective specific activities (**Figure 4 and Figure S4**). The triple-substituted variant NylCp2-TS^F134W/D304M/R330A^ (NylC-HP) achieved the highest performance exhibiting a 6.9-fold ± 0.3 improved specific activity for PA 6. Introducing a fourth substitution at position 99 (V6–V8), particularly D99R (V8), resulted in impaired activity (**Table S5**). What molecular interactions at this position affect polyamidase activity will be further evaluated in the computational analysis section of this study. Finally, the specific activity of NylC-HP could be enhanced from 77 ± 1.0 to 520 ± 19 µmol6-AHAeq. h^−1^ mgenzyme^−1^ in the first directed polyamidase evolution campaign. This brings the specific activity of NylC-HP within the range of the most efficient PETases (e.g., 963 µmolTPAeq. h^−1^ mgenzyme^−1^ for LCC^ICCG^) making NylC-HP the fastest polyamidase reported to this date.^1^

**Figure 4.**
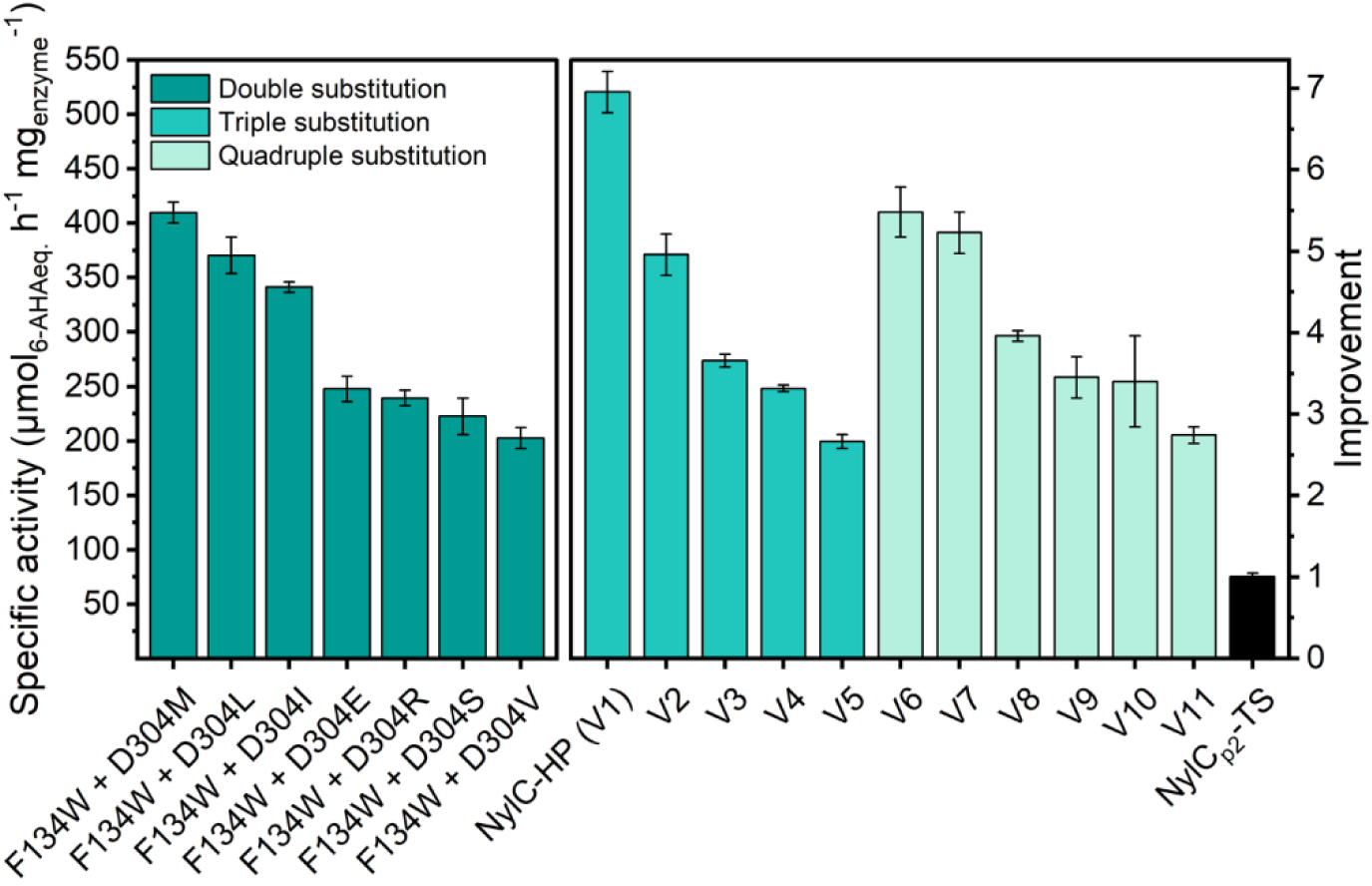
Specific activities of NylCp2-TS double-, triple-, and quadruple substitutions. Specific activities of double substitutions were calculated from released 6-AHA equivalents after incubation with Gf-PA 6 film (25 nM/0.95 mg L^−1^ enzyme, 456 g L^−1^ Gf-PA 6, 50 mM bicine, pH 8.0, 100 mM NaCl, 60 °C, 2 h). Specific activities of triple- and quadruple substitutions were calculated from the *V*max obtained from conventional Michaelis-Menten kinetics (25 nM/0.95 mg L^−1^ enzyme, 76–608 g L^−1^ Gf-PA 6, 50 mM bicine, pH 8.0, 100 mM NaCl, 60 °C, 2 h). Reactions were conducted in at least duplicate. Error bars represent the standard error of the mean.

### 2.2 Experimental characterization of NylC-HP

We optimized NylCp2-TS through directed evolution to enhance the PA 6 degradation efficiency and obtained a polyamidase with high performance harboring three substitutions (NylCp2-TS^F134W/D304M/R330A^; NylC-HP) (**Figure 5A**). In the following, we characterized NylC-HP experimentally.

**Figure 5.**
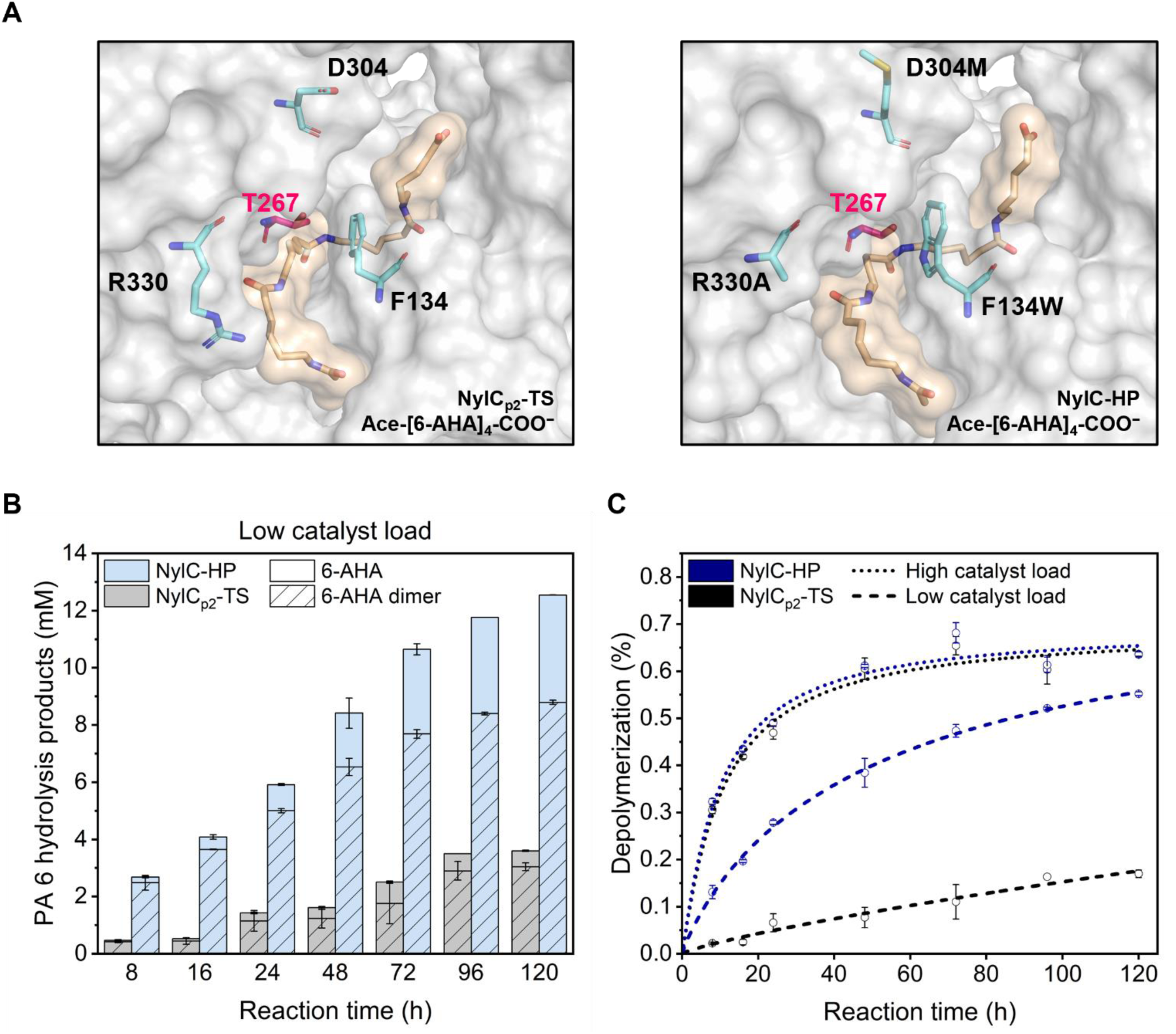
Experimental characterization of NylC-HP. A) Representative structure of the enzyme-substrate complexes NylCp2-TS/Ace-[6-AHA]_4_-COO^−^ and NylC-HP/Ace-[6-AHA]_4_-COO^−^(orientation 1). Introduced amino acid substitutions are shown as turquoise sticks. The catalytic threonine T267 is highlighted in pink. B) Time-resolved Gf-PA 6 degradation by NylCp2-TS and NylC-HP with low catalyst load (50 nM/1.90 mg L^−1^ enzyme, 456 g L^−1^ Gf-PA 6, 50 mM bicine, pH 8.0, 100 mM NaCl, 60 °C, 8–120 h). C) Degree of Gf-PA 6 depolymerization by NylCp2-TS and NylC-HP at low and high catalyst load. Reactions were conducted in at least duplicates. Error bars represent the standard error of the mean.

#### 2.2.1 NylC-HP preferentially degrades PA 6,6 over PA 6

We selected NylCp2-TS variants based on their PA 6 depolymerization efficiency with an up to 6.9-fold increased specific activity (**Figure 4 and Figure S4**). We also assessed the specific activity of NylC-HP towards PA 6,6 by detecting the release of the PA 6,6 monomer (6-[(6-Aminohexyl)amino]-6-oxo-hexanoic acid), the primary product of enzymatic PA 6,6 hydrolysis.^27^ Consistent with the results for PA 6, we observed a 3.4-fold ± 0.1 increase in specific activity towards PA 6,6 for NylC-HP (i.e., from 304 ± 9 µmolPA6,6monomer eq. h^−1^ mg ^−1^ to 1026 ± 25 µmolPA6,6monomer eq. h^−1^ mg ^−1^) compared to NylCp2-TS (**Figure S5**). Given that PA 6 hydrolysis was the primary selection pressure in our campaign, we anticipated the most significant improvement in specific activity towards PA 6. Interestingly, even though the achieved relative improvement of NylC-HP is higher for PA 6 hydrolysis (6.9-fold ± 0.3) compared to PA 6,6 (3.4-fold ± 0.1), the enzyme still hydrolyzes PA 6,6 two times faster than PA 6. This result is even more striking considering that NylC-HP does not fully hydrolyze PA 6,6 to hexamethylene diamine and adipic acid but stops at the PA 6,6 monomer (**Figure S6**). In contrast, PA 6 is at least partially converted to 6-AHA (**Figure 5B**).^27^ These observations emphasize that PA 6 and PA 6,6, despite sharing similar chemical and physical properties, undergo distinct enzymatic processing. A similar behavior was reported for the reaction kinetics of *Is*PETase for the semi-aromatic polymers PET and PEF, where the enzyme favored the furan-based PEF over the benzene-based PET.^37^

#### 2.2.2 NylC-HP depolymerizes PA 6 efficiently but limited degree of depolymerization

Parameters for the economic assessment of enzymatic polymer recycling processes have been recently defined as substrate loading, conversion/degree of depolymerization, and reaction kinetics, which together entail enzyme productivity.^1, 3^ To maximize space-time yields, substrate loading is best optimized by process engineering. For instance, polymers can be pre-treated to maximize substrate loading and surface of exchange, and minimize polymer crystallinity.^1, 28, 38^ In contrast, reaction kinetics and degree of depolymerization represent promising targets for protein engineers.^2, 39, 40^ Established metrics for assessing the performance of heterogeneous biocatalysts include monitoring the maximum reaction rate under substrate saturation/inverse Michaelis-Menten conditions (^inv^MM), which reveals the number of attack sites the depolymerase can effectively target; and under enzyme saturation/conventional Michaelis-Menten conditions (^conv^MM), which provides a measure of the maximum reaction rate of the depolymerization reaction.^41^ The degree of depolymerization can be displayed by ‘the ultimate mass’ fraction of depolymerized product but must be carefully distinguished from factors such as improved reaction kinetics, which can outperform polymer recrystallization and consequently increase the fraction of depolymerized polymer.^9^

To assess the kinetic efficiency of NylC-HP beyond initial rates, we conducted long-term degradation experiments under enzyme (low catalyst load) and substrate saturation (high catalyst load), determined the respective product composition, and subsequently computed the corresponding degree of depolymerization. At low catalyst load, NylC-HP exhibited a 5.8- fold ± 0.7 enhanced initial specific activity of 39.4 ± 4.2 mg6-AHAeq. h^−1^ mg ^−1^ within the first 8 hours of the reaction, compared to 6.8 ± 0.6 mg6-AHAeq. h^−1^ mgenzyme^−1^ for NylCp2-TS (**Figure ^5^B**). The average specific activity was 3.3-fold ± 0.1 higher with 11.1 ± 0.1 mg6-AHAeq. h^−1^ mg ^−1^ for NylC-HP and 3.4 ± 0.2 mg6-AHAeq. h^−1^ mg ^−1^ for NylCp2-TS throughout 120 h of reaction. At high catalyst load, NylC-HP and NylCp2-TS exhibited comparable initial and average specific activities of 97.2 ± 1.9 and 91.8 ± 2.4, and 12.7 ± 0.1 and 12.8 ± 0.1 mg6-AHAeq. h^−1^ mg ^−1^, respectively (**Figure S7**). The latter observation coincides with the final degree of depolymerization, where both enzymes stalled around 0.64 ± 0.01 % (w/w) (**Figure 5C**). This highlights that increasing the amount of targetable attack sites and the degree of depolymerization is crucial to unlock the potential of polyamidases for PA recycling.

#### 2.2.3 NylC-HP shows higher conversion to the PA 6 monomer than NylCp2-TS

The enzymatic depolymerization of PA 6 is complex, involving the hydrolysis of distinct attack sites on the polymer surface and oligomers with different chain lengths. This makes determining reaction rates for individual partial reactions challenging.^27^ The two main products of PA 6 hydrolysis are 6-AHA and the corresponding dimer. Tuning the conversion towards 6-AHA is desirable since, in a circular process, 6-AHA can be converted to ε-caprolactam and repolymerized to virgin-grade PA 6.^42^ After 120 hours, NylC-HP achieved notably higher rates of complete depolymerization compared to the parental enzyme. Specifically, 30 % (low catalyst load) and 90 % (high catalyst load) of the released products were fully depolymerized to 6-AHA, whereas the parental enzyme achieved only 15 % and 80 %, respectively (**Figure 5B and Figure S7**).

#### 2.2.4 NylC-HP possesses enhanced thermal stability

Next to depolymerase activity, thermal stability plays a pivotal role in enzymatic polymer recycling, enabling accelerated reaction rates at elevated temperatures and reducing catalyst deactivation during the process.^43, 44^ Consequently, we determined melting temperatures for all tested variants (**Figure 6**). Substitutions at positions D99, F301, and R330 generally reduced thermal stability, except for a slight increase for D99R. The substitution F134W had no impact on thermal stability. Interestingly, all single-substitutions at D304 except D304W increased thermal stability by up to 3.8 ± 0.1 °C and for all NylCp2-TS^F134W/D304X^ double-substitutions by up to 4.8 ± 0.2 °C. Although NylCp2-TS is already thermostable (*T*m = 85.7 ± 0.1 °C), we were able to further improve the thermal stability of NylC-HP by an additional 4.2 ± 0.2 °C, achieving a remarkable *T*m of 89.9 ± 0.1 °C.

**Figure 6.**
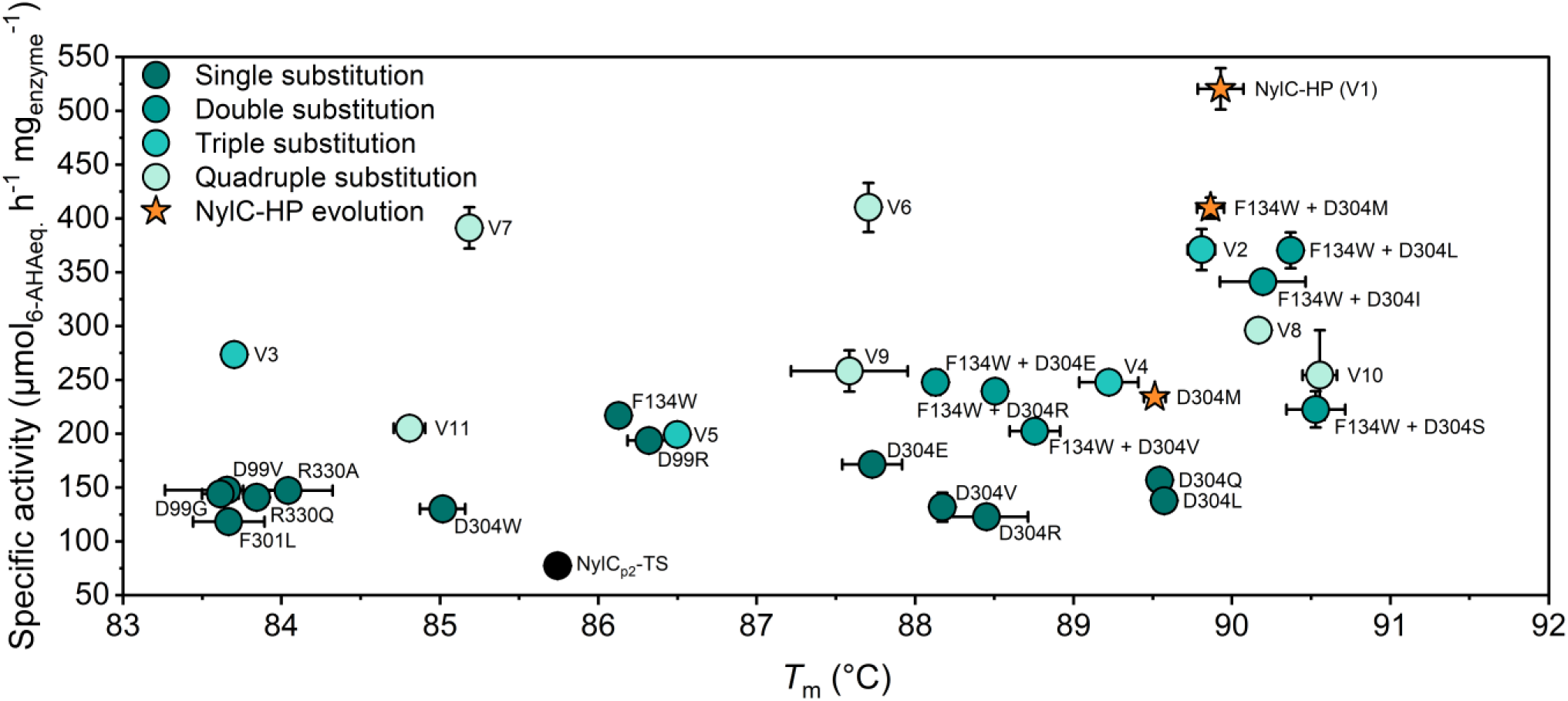
Thermal stability and specific activity of NylCp2-TS variants. Melting temperatures (*T*m) of NylCp2-TS variants are plotted against their specific activity. The parental enzyme, NylCp2-TS, is shown in black, with variants of increasing degrees of substitution represented in progressively brighter turquoise tones, and the evolutionary trajectory of NylC-HP highlighted in orange. Reactions were conducted in at least duplicates. Error bars represent the standard error of the mean.<

The enhanced specific activity underlines NylC-HP’s PA 6 depolymerization efficiency, leading to enhanced conversion rates and an increased monomer yield. The degree of depolymerization, however, remains a limiting factor, as previously reported.^10, 12^ Regardless, the development of NylC-HP represents hitherto unmatched enzymatic PA 6 and PA 6,6 depolymerization efficiency together with excellent thermal stability, rendering NylC-HP not only the fastest but also one of the most thermostable characterized polyamidases to this date.

### 2.3 Elucidating structure-function relationships using docking and MD simulations

To gain insights into molecular changes of enzyme-substrate interactions induced by the most beneficial substitutions D99R, F134W, D304M, and R330A, we generated enzyme-substrate complexes, using NylCp2-TS, NylC-HP, and NylC-HP^D99R^ as enzymes and Ace-[6-AHA]4-COO^−^ and NMe-[6-AHA]4-NH3^+^ as substrates. We applied an incremental docking procedure (**Figure S8-S9**), followed by MD simulations (**Figure S10-S12**) and compared the dynamic behavior of examined enzyme-substrate complexes. Moreover, we studied the dynamics of systems containing – besides water and ions – only the enzyme (i.e., NylCp2-TS, NylCp2-TS^D99R^, NylC-HP, and NylC-HP^D99R^) to determine differences in protein-protein interactions (**Figure S10-S13**). In addition to the identification of activity-enhancing molecular changes, deciphering the driving force that causes the negative epistasis that was observed upon combination of D99R and D304M, and structural features limiting the degree of depolymerization was of particular interest.

#### 2.3.1 Improved variants NylC-HP and NylC-HP^D99R^ prefer different substrate orientations than NylCp2-TS

The incremental docking procedure revealed differences in the substrate poses that were ranked best (high population and affinity) for each enzyme. While for NylCp2-TS both orientations of the substrate Ace-[6-AHA]4-COO^−^ led to the energetically most favored conformations, the best substrate poses for NylC-HP and NylC-HP^D99R^ could be achieved with Ace-[6-AHA]4-COO^−^ (orientation 1) and NMe-[6-AHA]4-NH ^+^ (orientation 2) (**Figure S2, Table S3, and Table S7-S8**).

#### 2.3.2 D99 and R330 stabilize substrate termini through electrostatic interactions

During simulation we observed that the residues at positions D99 and R330 interact with the charged termini of the polymer, COO^−^ and NH_3_ ^+^ (**Figure 7A and Figure 8A**). At least one hydrogen bond (H-bond) or salt bridge between the side chain of R330 and the negatively charged terminus of the substrate was formed for 67 ± 16 % (H-bond) and 65 ± 18 % (salt bridge) of the simulation time, respectively (**Figure 7A and Table S9**). In addition, we found a significant drop in interaction energy (sum of short-range Lennard-Jones and Coulomb interaction energy) between the enzyme and the substrate upon H-bond formation with R330 Δ*E* = −98 ± 18 kJ mol^−1^ (**Table S10**). Variants comprising the substitution R330A showed a significantly reduced contact frequency (*f*) between the residue at this position and the substrate compared to NylCp2-TS (Δ*f*(NylC-HP, NylCp2-TS) = −51 ± 11 % and Δ*f*(NylC-HP^D99R^, NylCp2-TS) = −47 ± 11 %). Depolymerases tend to exhibit excessively strong binding to their polymer substrates, and reducing binding affinity, has been shown to enhance activity.^45^ As observed during MD simulations, exchanging arginine with alanine reduces attractive electrostatic interactions between enzyme and substrate, which may explain the increased specific activity observed upon introducing the R330A substitution.

**Figure 7.**
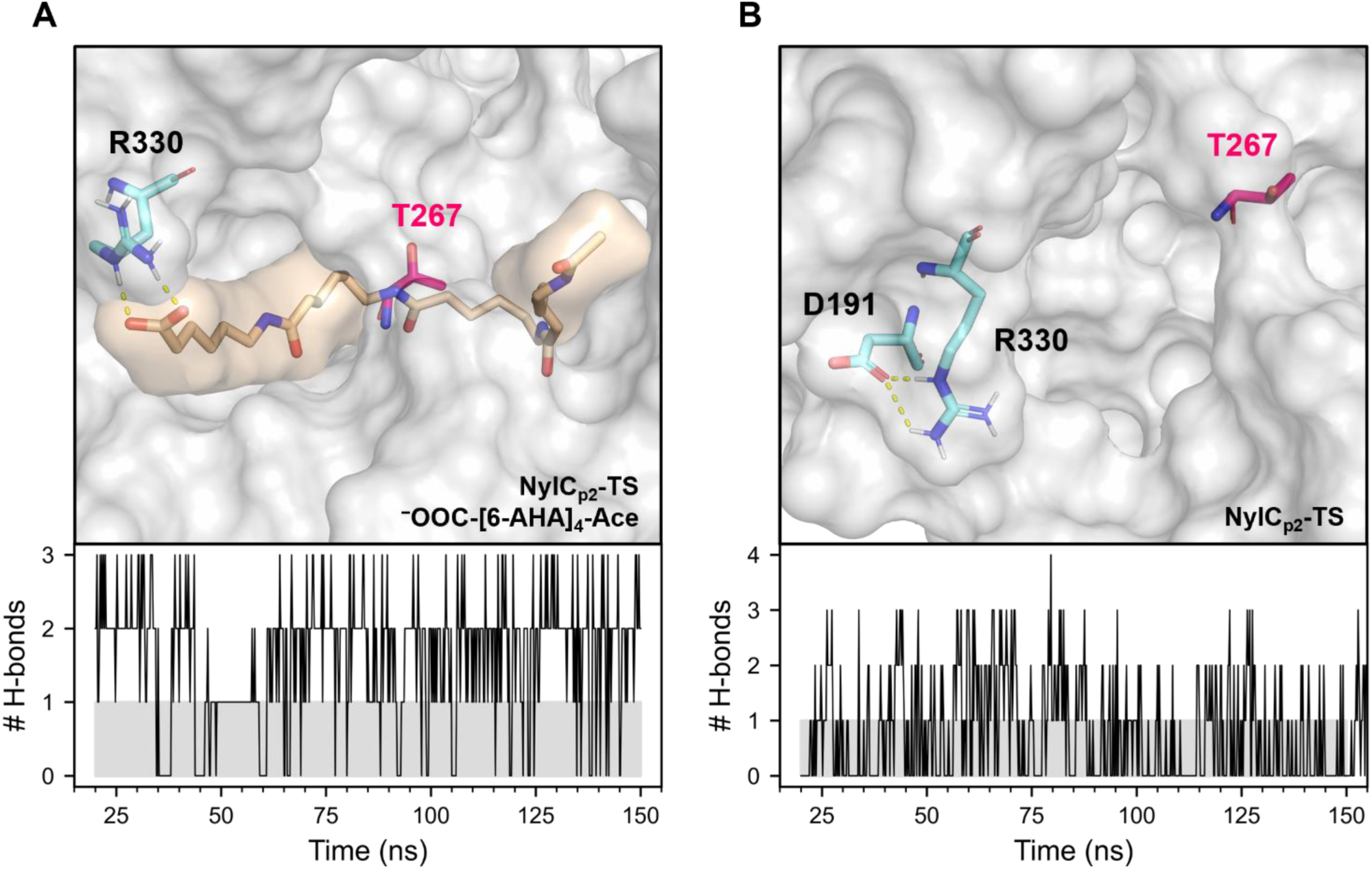
Hydrogen bonds (H-bonds) of residue R330 observed during molecular dynamics (MD) simulations. A) Representative structure of the enzyme-substrate complex NylCp2-TS/^−^OOC-[6-AHA]4-Ace with established H-bonds between the side chain of residue R330 and the negatively charged terminus of the substrate (top) and the number of such bonds (# H-bonds) in dependency of the simulation time observed for one MD simulation of this system (bottom). B) Representative structure of NylCp2-TS with established H-bonds between the side chains of residues R330 and D191 (top) and the number of such bonds (# H-bonds) in dependency of the simulation time observed for one MD simulation of this system (bottom).

**Figure 8.**
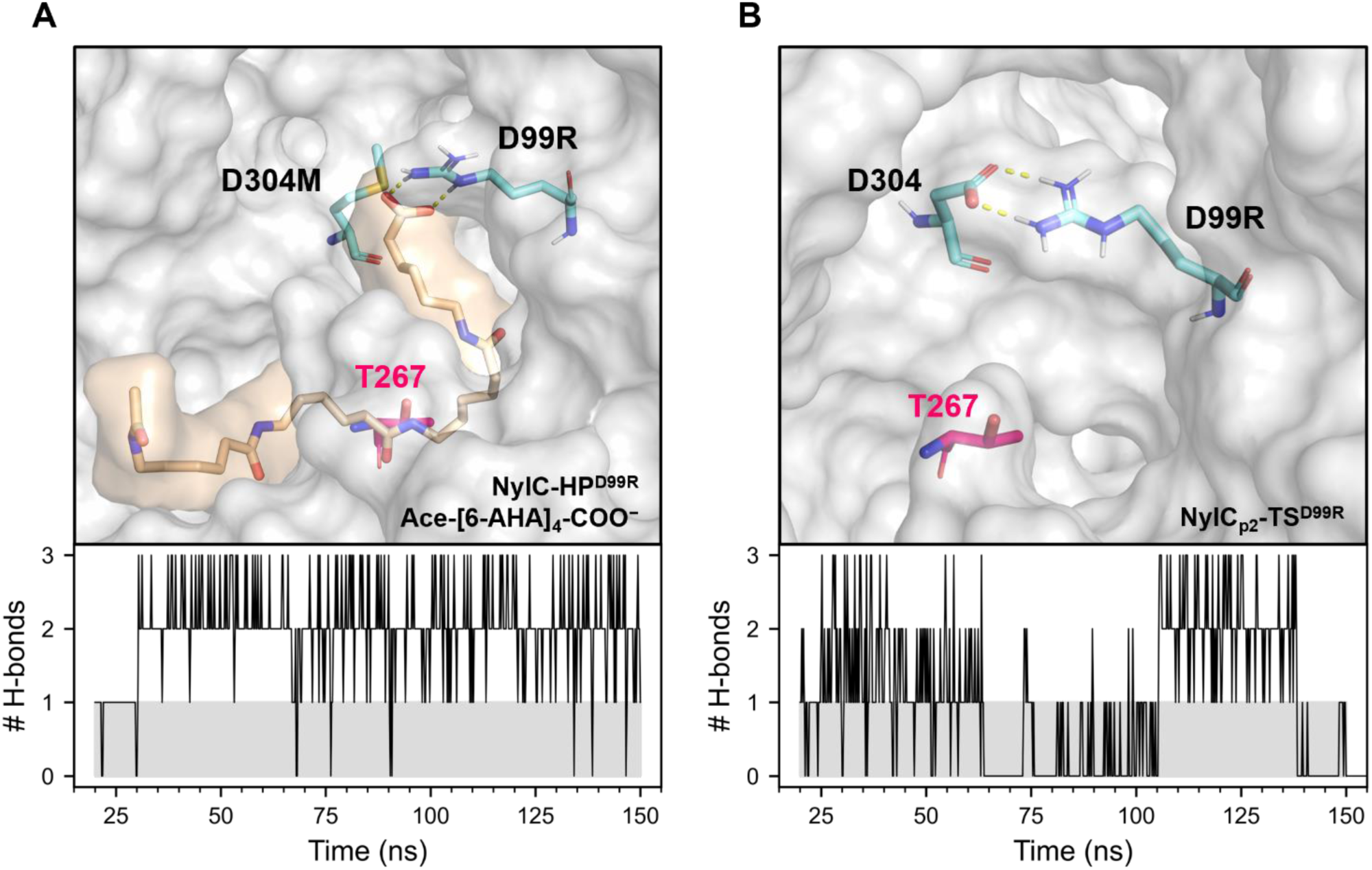
Hydrogen bonds (H-bonds) of residue D99R observed during molecular dynamics (MD) simulations. A) Representative structure of the enzyme-substrate complex NylCp2-TS^D99R^/Ace-[6-AHA]4-COO^−^ with established H-bonds between the side chain of residue R99 and the negatively charged terminus of the substrate (top) and the number of such bonds (# H-bonds) in dependency of the simulation time observed for one MD simulation of this system (bottom). B) Representative structure of NylCp2-TS^D99R^ with established H-bonds between the side chains of residues R99 and D304 (top) and the number of such bonds (# H-bonds) in dependency of the simulation time observed for one MD simulation of this system (bottom).

In addition to altered enzyme-substrate interactions, we also observed differences in protein-protein interactions, which manifested itself in the dissolution of the salt bridge between the residues D191 and R330 (Figure 7B). While the formation of such a salt bridge could be observed up to 28 % of the simulation time for the NylCp2-TS variant, it was absent for all variants carrying the substitution R330A (Table S11). It has been shown that the (thermal) stability of a protein strongly correlates with the amount of non-covalent interactions present in the structure (e.g., H-bonds, salt bridges, and hydrophobic interactions).^46–48^ We propose that the measured reduced thermal stability of NylCp2-TS^R330A^ compared to NylCp2-TS may be explained by the omission of the structure-stabilizing interaction D191-R330 when introducing the substitution R330A.

The increase in specific activity of NylC-HP was achieved by substituting F134W, D304M, and R330A (**Figure 4 and Table S5**). Introducing D99R to NylCp2-TS also increased enzyme activity, however, when introduced to NylC-HP, negative epistasis was observed. Specifically, the improvement in specific activity decreased from 6.9- to 3.9-fold for NylC-HP and NylC-HP^D99R^. For NylC-HP^D99R^, the MD simulations revealed persistent H-bond formation (up to 98 % of the simulation time) between D99R and the negatively charged terminus of the substrate Ace-[6-AHA]4-COO^−^ (**Figure 8A and Table S9**). We determined the interaction energy between the enzyme and the substrate for this system and observed a significant drop of Δ*E* = −92 ± 26 kJ mol^−1^ upon H-bond formation between D99R and the carboxy-terminus of the substrate (**Table S10**). For NylCp2-TS^D99R^, we observed the formation of a salt bridge between D99R and D304 during simulation (39 ± 10 % of the simulation time) (**Figure 8B and Table S11**). This salt bridge might also explain the increase in *T*m observed for NylCp2-TS^D99R^ (**Figure 6**).

Based on the MD results described above, we assume that reducing interactions between residue 99 and the charged terminus of the polymer supports substrate hydrolysis. In the case of NylCp2-TS^D99R^, substrate interactions may be reduced, as salt bridge formation between the guanidino group of D99R and the carboxy group of D304 could prevent D99R from interacting with the substrate. However, when residue 304 cannot serve as a H-bond acceptor, as in NylC-HP^D99R^, the interaction between D99R and the substrate may be restored, leading to a decrease in enzyme activity.

#### 2.3.3 Positions 304 and 305 are non-conserved islands in a highly conserved region

The strong beneficial impact on NylCp2-TS activity and thermal stability of all tested D304X substitutions suggests that the naturally occurring aspartate causes a structurally unfavorable conformation of the enzyme. Conservation analysis showed that positions D304 and M305 constitute the sole highly variable positions within a predominantly conserved patch spanning residues of the catalytic triad (i.e., D306 and D308) (**Figure 9**). Interestingly, in contrast to saturating position D304, saturating position M305 led to no variant with notable polyamidase activity.

**Figure 9.**
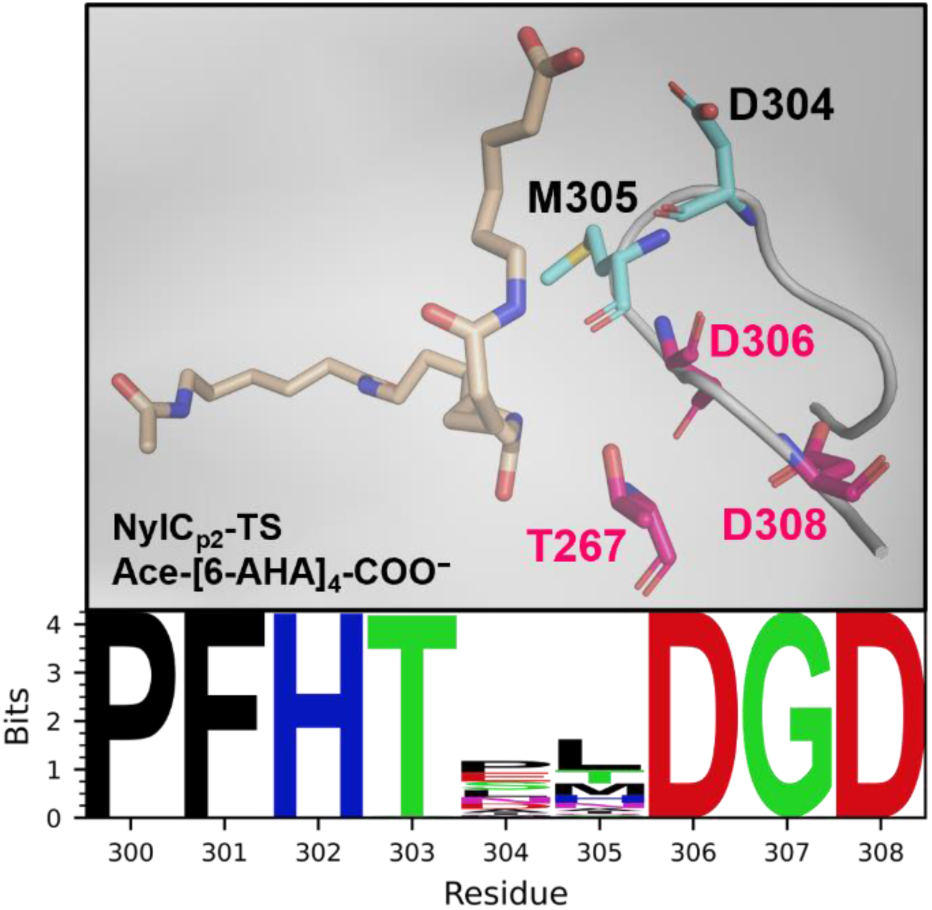
Residues D304 and M305 represent non-conserved amino acids within a highly conserved region of NylCp2-TS that includes two residues of the catalytic triad (i.e., D306 and D308). Detailed view of the highly conserved region including positions 300-308 of NylCp2-TS, with the non-conserved residues highlighted in blue and the residues of the catalytic triad highlighted in pink (top). Excerpt from the sequence logo of NylCp2-TS (bottom).

#### 2.3.4 Conformational dynamics of aromatic sidechains at position 134 control substrate binding and hydrolysis

While introducing various amino acids at positions D99, D304, and R330 led to more active enzyme variants, F134W was the only substitution that improved enzyme activity at position F134 (**Figure 3**). The MD simulations revealed that the aromatic side chain of the residue at position F134 mainly occurs in two conformations: (I) vertical, pointing towards the active site (**Figure 10A and Table S12-S15**) and (II) horizontal or vertical, pointing out of the enzyme (**Figure 10B and Table S12-S15**). We observed that the first conformation fixes the amide bond near the nucleophilic residue T267 and prevents the substrate from establishing intramolecular H-bonds that causes the polymer chain to fold. Additionally, for variants harboring the F134W substitution, H-bonds between the substrate and the indole side chain were established for up to 25 % of the simulation time (**Table S16**). Thereby, F134W can fix the amide bond of the substrate near the active residue T267 which may support substrate hydrolysis. Conformation 2 increases the free volume inside the binding pocket of the enzyme, allowing the substrate to move more freely and establish intramolecular H-bonds. We hypothesize that conformation 1 supports substrate hydrolysis and conformation 2 is responsible for substrate binding. As both conformations are essential for polymer degradation, it may explain why only a conservative substitution to an aromatic residue, which is similar in volume and shape to the wild-type amino acid phenylalanine and thus can realize both conformations, leads to an active enzyme variant. Interestingly, a phenylalanine-to-tyrosine substitution was not observed during screening.

**Figure 10.**
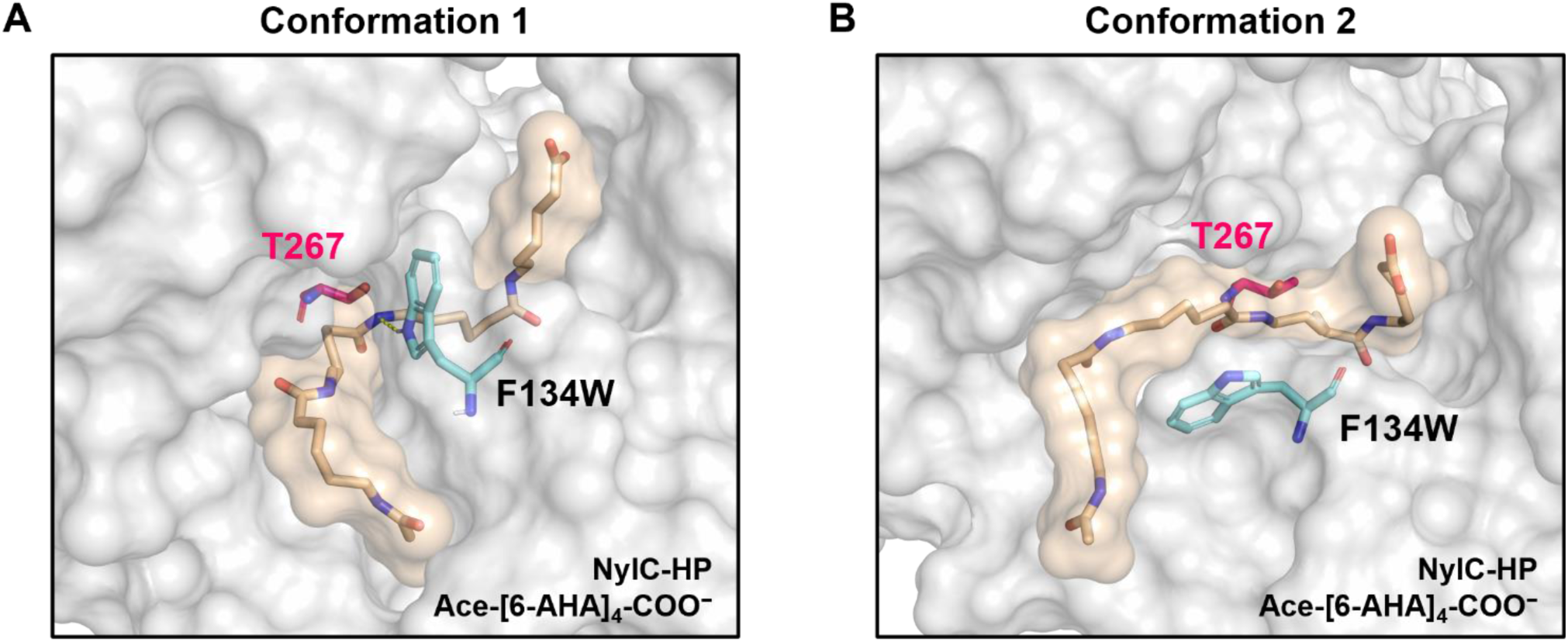
Representative structures highlighting the two main conformations of residue F134W observed during molecular dynamics simulations of the enzyme-substrate complex NylC-HP/Ace-[6-AHA]4-COO^−^. A) Conformation 1 is characterized by a vertically oriented side chain of F134W, which prevents intramolecular interactions and (optionally) fixes the amide bond to be cleaved by H-bonding close to the active residue T267. B) Conformation 2 is characterized by a horizontal alignment of the side chain of F134W. This generates free space in the binding pocket and intramolecular interactions of the substrate can be established.

#### 2.3.5 Incremental substrate docking and MD simulations provide insights into PA 6 hydrolysis and the structural limitations of NylC-type enzymes

Based on the computational analysis, we attribute the positive effects on enzyme activity predominantly to the (I) reduction of steric and electrostatic interactions with the substrate, realized by the substitutions D99R and R330A, (II) elimination of unfavorable interactions, implemented by substitutions at position D304, and (III) formation of a H-bond that fixes the amide bond near the active residue T267 by introducing F134W. In addition, the residue at position F134 appears to be crucial for opening/closing the binding pocket and preventing the substrate from establishing intramolecular interactions. The negative epistasis observed upon combining D99R with a substitution at position D304 is likely a result of the restoration of unfavorable enzyme-substrate interactions. The combination of incremental docking and subsequent MD simulations enabled targeted and efficient enzyme engineering with large substrates. Moreover, MD simulations could be applied to generate understanding of changes in intra- (i.e., protein-protein and substrate-substrate) and intermolecular (i.e., enzyme-substrate) interactions caused by substitutions that affected enzyme activity and stability.

The KnowVolution campaign yielded a polyamidase, NylC-HP, with the highest specific activity without improving the degree of depolymerization. Recently a promising strategy was presented where the degree of depolymerization of PETases could be enhanced.^2^ By increasing the flexibility of the PET-binding groove of BhrPETase, it was possible to yield TurboPETase, which operates on versatile surface structures through dynamic binding. Restructuring of the active site broadened the spectrum of attack sites accessible for enzymatic hydrolyzation and led to enzymes that nearly completely depolymerize PET at industrially relevant solids loading.^2, 49^ Our simulations revealed that the active site of NylC-HP is located in a narrow pocket, which only offers space for a single PA 6 chain (**Figure 5A**). We hypothesize that NylC-HP can only hydrolyze exposed polymer chains, which explains the low degree of depolymerization (**Figure 5C**). To achieve a higher degree of depolymerization, the enzyme would need to strip PA chains from the polymer surface and break their intra- and interchain interactions, before substrate binding to the relatively narrow binding pocket and subsequent hydrolysis can occur. Considering the low degree of depolymerization achieved, NylC-HP appears to be incapable of stripping off PA 6 chains that are firmly attached to the polymer. Finally, NylC-HP can cause surface alterations but cannot induce pitting and penetrate the inner polymer as observed for PETases.^10, 12, 37, 50^ In comparison to TurboPETase, the active site of NylC-HP is more buried inside the enzyme. Therefore, a comprehensive redesign and substantial widening of the binding pocket in NylC-HP would be required to enhance the degree of PA 6 and PA 6,6 depolymerization.

## 3 Conclusion

Enzymatic recycling of synthetic polymers has emerged as a promising alternative to traditional recycling methods. However, environmentally friendly and cost-effective polymer hydrolysis processes rely on stable enzymes with high activity and high degree of depolymerization. We engineered NylCp2-TS and developed the highest-performing polyamidase to date, NylC-HP. Through the engineering of NylCp2-TS and in-depth MD simulations of enzyme-substrate interactions, we identified key positions (D99, F134, D304, and R330) crucial for substrate interactions and hydrolysis, ultimately resulting in a 6.9-fold improved *k*cat. Our study shows that the combination of incremental docking and molecular dynamics simulation is an excellent guide for the efficient engineering of polyamidases with high specific activity, while simultaneously generating molecular understanding. The incremental approach is particularly effective for engineering enzymes to process flexible polymers with numerous rotatable bonds, such as linear polyamides, -esters, and -urethanes. Interestingly, active site engineering did not result in any improvements of the degree of depolymerization. We conclude that large parts of PA 6 and PA 6,6 are not accessible for hydrolysis by NylC-HP which could be attributed to an insufficient binding pocket size and/or flexibility. We anticipate that upcoming engineering campaigns will generate valuable insights, leading to strategies for remodeling binding pockets to enhance the degree of depolymerization for polyamidases. Moreover, novel polyamidases will be identified which might represent a more promising starting point for engineering with a naturally broader active site. These can then be subjected to the general computational approach outlined in this study to enhance their activity.

## 4 Experimental section

### 4.1 Materials

Gf-PA 6 film (0.2 mm thick, AM30-FM-000200) and Gf-PA 6,6 granules (3 mm in diameter, AM32-GL-000115) were acquired from Goodfellow GmbH (Hamburg, Germany). For the Gf-PA 6 film, 6.4 mm diameter discs (7.6 mg each) were punched using a 12 kN 96-well puncher (Gechter, Germany).

### 4.2 Experimental methods

#### 4.2.1 Gene construction

The gene encoding the parental enzyme, NylCp2-TS, was generated by the commercial synthesis of the NylCp2 wildtype by Eurofins, cloning into pET 21a (+) (Novagen), and introduction of four amino acid substitutions to obtain the thermostable NylC^D36A/D122G/H130Y/E263Q^ (NylCp2-TS).^16^

#### 4.2.2 Library construction

Site-saturation mutagenesis (SSM) libraries of NylCp2-TS were generated using NNK primers (**Table S17**). Polymerase chain reaction (PCR) was conducted using Q5^®^ High-Fidelity DNA Polymerase (New England BioLabs, USA). PCR products were incubated with DpnI (New England BioLabs, USA; 37 °C, 60 min) to digest template DNA, purified, and transformed into chemically competent *E. coli* BL21(DE3) cells.

#### 4.2.3 Library cultivation

SSM libraries were cultivated in 96-well F-bottom MTPs (Greiner, Frickenhausen, Germany) by inoculating 200 µL lysogeny broth media (LB media, 1 % tryptone, 0.5 % yeast extract, and 1 % NaCl, 50 µg mL^−1^ carbenicillin) with a single freshly transformed colony per well. 12 wells were inoculated with NylCp2-TS as parental enzyme reference. After overnight incubation (900 RPM, 37 °C, 70 % humidity, 16 h), 10 µL pre-culture were added to 130 µL Terrific broth media (TB media, 24 g L^−1^ yeast extract, 20 g L^−1^ tryptone, 4 mL L^−1^ glycerol, 0.017 M KH2PO4, 0.072 M K2HPO, pH 7.4, 50 µg mL^−1^ carbenicillin) in 96-well V-bottom MTPs. Following growth to an OD600 of 0.6 (900 RPM, 37 °C, 70 % humidity, 2 h), expression was induced by the addition of 10 µL isopropyl-*β*-D-thiogalactopyranoside (IPTG, 1.5 mM). Cultures were further incubated overnight (900 RPM, 20 °C, 70 % humidity, 20 h) and harvested by centrifugation (Eppendorf 5810R; 3,220 x *g*, 4 °C, 15 min). The obtained cell pellet was resuspended with 150 µL buffer (50 mM Bicine, pH 8.0, 100 mM NaCl, 20 °C, 10 min) to remove the residual medium and pelleted again. Cells were disrupted by resuspension and incubation in 150 µL lysis buffer (50 mM Bicine, pH 8.0, 1.5 mg mL^−1^ lysozyme, 25 µg mL^−1^ DNAseI; 900 RPM, 37 °C, 2 h). Insoluble cell residue was pelleted (Eppendorf 5810R; 3,220 x *g*, 4 °C, 15 min) and the cell-free extract was used for the screening.

#### 4.2.4 Library screening

Screening for improved enzyme variants was conducted using the AMIDE method.^12^ Screening reactions were initiated by adding 10 µL of clarified cell lysate to 190 µL screening buffer (50 mM Bicine, pH 9.0, 100 mM NaCl) containing 38 g L^−1^ Gf-PA 6 film in a 96-well F-bottom MTP. Before incubation (800 RPM, 60 °C, 4 h), plates were sealed (EasySeal™, Greiner GmbH, Germany) to prevent evaporation. Plates were then centrifuged to precipitate denatured protein (Eppendorf 5810R, 4 °C, 3,220 x *g*, 20 min), and 100 µL of the reaction mixture was analyzed via MAFC assay.

#### 4.2.5 MAFC assay

Released amines were detected by MAFC MTP assay. To prepare the MAFC assay solution, 4 mM Meldrum’s acid furfural conjugate (MAFC) were dissolved in 100 % ethanol and filtered through a 0.45 µm syringe filter (Aerodiscs®, Pall Deutschland Holding GmbH & Co. KG, Germany). Next, plates were developed (800 RPM, 37 °C, 30 min), centrifuged (Eppendorf 5810R, 3,220 x *g*, 4 °C, 1 min), and the absorbance was measured at 494 nm (CLARIOstar, BMG LABTECH, Germany; 100 flashes/well).

#### 4.2.6 High-performance liquid chromatography (HPLC) analysis of PA degradation products

PA 6 and PA 6,6 degradation products (6-aminohexanoic acid, adipic acid, PA 6 dimer, and PA 6,6 dimer) were analyzed using a Shimadzu prominence LC system together with a C18-RP column (NUCLEOSIL 100-5 C18, Macherey-Nagel GmbH & Co. KG, Germany). Analytes were quantified via a photodiode array (Nexera X2 SPD-M30A, Shimadzu, Japan) at 220 nm. Mobile phase A comprised 0.1 % sulfuric acid and mobile phase B comprised 100 % acetonitrile. Analyte injections of 10 µL were isocratically resolved. For PA 6, 5 % mobile phase B with a constant flow rate of 1 mL min^−1^ over a total runtime of 20 min was applied. For PA 6,6 15 % mobile phase B with a constant flow rate of 1 mL min^−1^ over a total runtime of 20 min was applied.

#### 4.2.7 Michaelis-Menten kinetics

To determine the maximum reaction velocity, enzyme activity was assayed at enzyme saturation (Gf-PA 6 film, 38–684 g L^−1^; Gf-PA 6,6 granules, 110–880 g L^−1^), and the curves were fitted using the Hill equation (**Equation 1**) with *v* being the enzymatic reaction rate observed at a particular substrate concentration, *V*max the maximum reaction velocity, [S] the substrate concentration, *K* the dissociation constant, representing the substrate concentration at which half of the binding sites are occupied, and *n = 2* as fixed Hill coefficient using OriginPro® 2024 (OriginLab). Kinetic parameters were derived from the Michaelis-Menten Equation (**Equation 2**), with *v* being the enzymatic reaction rate observed at a particular substrate concentration, *V*max the maximum reaction velocity, [S] the substrate concentration, and *K*M the Michaelis-Menten constant.

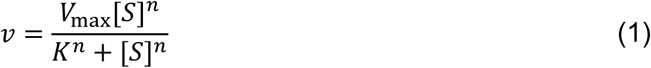

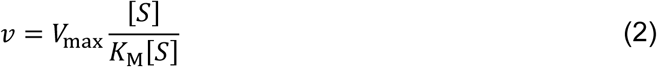

#### 4.2.8 Chemical synthesis of MAFC

MAFC was synthesized as previously reported.^51^ Briefly, 1.51 g (10.5 mmol) of 2,2-dimethyl-1,3-dioxane-4,6-dione was mixed with 961 mg (10 mmol) of 2-furaldehyde in 30 mL water and stirred at 70 °C for 2 h. The product precipitates as yellow solid and was dissolved in dichloromethane, washed with saturated aqueous NaHSO3, H2O, NaHCO3, and brine (30 mL each), and filtered. Finally, organic solvents were evaporated to obtain a bright yellow powder.

### 4.3 Computational methods

#### 4.3.1 Incremental docking Enzyme (receptor) preparation

First, the structure of the 6-aminohexanoate-oligomer hydrolase NylCp2-TS (PDB identifier 5Y0M) was fetched from the RCSB Protein Data Bank.^52, 53^ Heteroatoms were removed and symmetry mates were generated to obtain the tetrameric structure using PyMOL™ Molecular Graphics System (version 2.3.0, Schrödinger, LLC). The missing residues V29 and F30 were modeled with MODELLER (version 10.4).^54^ During this procedure the non-missing residues were fixed. Further, MODELLER was used to generate enzyme variants by introducing the variant-specific substitutions. The N-terminal residues D19 and I16 were capped with an acetyl group (Ace), while an N-methyl (NMe) cap was used for the C-terminal residues P259 and P260. The protonation state of the enzyme was adjusted according to a pH value of 8.0.

Energy minimization was performed with GROMACS (version 2022.3) using the CHARMM36 force field (version July 2022) and the TIP3P water model.^55–57^ For energy minimization, the system was solvated, neutralized, and a concentration of 100 mM NaCl was adjusted. We applied the „split the charge difference in two“ rule to add proper amounts of ions.^58^ For energy minimization, the steepest descent algorithm was applied until a maximum force of 1000 kJ mol^−1^ nm^−1^ on any atom was reached. The energy-minimized enzyme structure was extracted and receptor preparation was performed with AutoDockFR (version 1.2rc1).^59^ The substituted residues at positions D99, F134, D304, and R330 were treated as flexible during the docking procedure.

##### Substrate (ligand) preparation

We used Ace-[6-AHA]4-COO^−^ and NMe-[6-AHA]4-NH3^+^ oligomers as model substrates to mimic the PA 6 chain and study the enzyme-substrate interactions (**Figure 1**).

The substrates (fragments) were generated and energy minimized with AVOGADRO (version 1.2.0).^60^ Substrate preparation was performed with AutoDockFR (version 1.2rc1).^59^

##### Incremental docking procedure

To overcome the sampling problem when performing docking experiments with substrates that have many rotatable bonds, we implemented and applied a parallelized, incremental meta-docking procedure inspired by DINC. ^24, 61, 62^ Our workflow can be summarized as follows:

1. Select Ace-[6-AHA]-NMe as the first fragment to be docked to the prepared enzyme structure.
2. To ensure broad conformation sampling and efficient use of computing resources, we performed 240 parallel runs of AutoDock Vina (version 1.2.5) to generate a pool of fragment poses.^63, 64^ The generated conformations were clustered based on their root mean square deviation using a cutoff of 1.5 nm and structures that presented the bond to be cleaved near the active residue T267 were selected for the next round.
3. The selected fragments of the previous round were extended by another unit of 6-AHA and the bonds of the parental fragment were set to be non-rotatable. The structure was energy minimized with AVOGADRO (version 1.2.0) and prepared for docking with AutoDockFR (version 1.2rc1).^59^ After extension and preparation, conformation sampling was executed as described in 2.
4. Iterative cycles of 2 and 3 were performed until the structure consisted of four units of 6-AHA.To avoid bias, we performed fragment extension such that both orientations of each substrate (i.e., Ace-[6-AHA]4-COO^−^ and NMe-[6-AHA]4-NH ^+^) were generated (**Figure S8-S9**). The final poses were clustered and evaluated based on their affinity and population. Poses that had a high population and a low free energy of binding were selected for molecular dynamics (MD) simulation.

#### 4.3.2 Molecular dynamics simulation

MD simulations were performed to study the dynamic behavior of different enzyme variants and enzyme-substrate complexes. At first, parameters were assigned to the substrates using the CHARMM general force field program (version 2.5).^65, 66^ MD simulations were performed with GROMACS (version 2022.3) using the CHARMM36 force field (version July 2022) and the TIP3P water model.^55–57^ Hydrogens were added to the side chains to mimic a pH value of 8.0. The enzyme(-substrate complex) was centered in a cubic box with 1 nm between the solute and the box. The system was solvated, neutralized, and a concentration of 100 mM NaCl was adjusted. To ensure neutralization while not exceeding the concentration of 100 mM NaCl, we applied the „split the charge difference in two“ rule.^58^ Energy minimization was performed applying the steepest descent algorithm until a maximum force of 1000 kJ mol^−1^ nm^−1^ on any atom was reached. The system was equilibrated by a 1 ns *NVT* run at 293.15 K using the modified Berendsen thermostat (velocity rescaling), while coupling protein(/substrate) and water/ions separately to the temperature bath using a coupling constant of 0.1 ps, followed by a 1 ns *NpT* run at 293.15 K and 1.013 bar using the previously described modified Berendsen thermostat and the Parinello-Rahman barostat with the following parameters: pcoupltype = isotropic, tau_p = 1.0 ps, and compressibility = 4.5 · 10^−5^. For equilibration, position restraints were applied to the protein (and the substrate) using the linear constraint solver (LINCS) algorithm. To generate simulation data, the position restraints were removed, and a 150 ns production run was performed under *NpT* conditions for the enzyme-substrate complexes and a 200 ns run was performed for the enzyme-only systems. The leap-frog algorithm was used with an integration time of 2 fs for solving Newton‘s equations of motion. Particle-Mesh Ewald electrostatics were used to calculate the Coulomb interactions with a short-range electrostatic and a van der Waals cutoff of 1.0 nm. Three independent MD runs were performed for each system. The first 20 ns of the simulation were not considered for analysis (**Figure S10-S13**).

#### 4.3.3 Conservation analysis

First, a protein-protein BLAST was run to extract a set of similar sequences from the non-redundant protein sequences database (update date 2024/06/10) using the sequence of NylCp2-TS (PDB identifier 5Y0M) as query.^52, 67^ The maximum number of aligned sequences to display was increased to 5000, while the other algorithm parameters remained unchanged. Next, a multiple sequence alignment was constructed with Clustal Omega (clustalo version 1.2.4) using sequences with an *E*-value ≤ 1 · 10^−100^ and a sequence identity ≤ 95 %.^68–70^ The sequence logo was generated according to the procedure described by Schneider et al.^71^

## Acknowledgements

Hendrik Puetz and Alexander-Maurice Illig contributed equally to this work.

We gratefully thank Prof. Maria Fyta, PhD and Daniela Paula Herrera Toro, PhD for the lively discussions and their valuable suggestions on docking and molecular dynamics simulations.

We would like to thank Dr. Marian Bienstein for his support in applying for computing time. Computations were performed with computing resources granted by RWTH Aachen University under project rwth1584.

We thank the German Federal Ministry for Education and Research for funding our study via Robuste und selektive lipolytische Biokatalysatoren für industrielle Anwendungen (LipoBioCat I and II) [BMBF Project FKZ 031B0837C, and O31B1342C].

## Author contribution

**Hendrik Puetz:** Conceptualization, data curation, formal analysis, funding acquisition, investigation, methodology, project administration, supervision, visualization, writing – original draft, writing – review & editing.

**Alexander-Maurice Illig:** Conceptualization, data curation, formal analysis, investigation, methodology, software, visualization, writing – original draft, writing – review & editing.

**Mariia Vorobii:** Investigation, writing – review & editing.

**Christoph Janknecht:** Investigation.

**Francisca Contreras Leiva:** Funding acquisition, supervision, writing – review & editing.

**Fabian Flemig:** Investigation.

**Ulrich Schwaneberg:** Funding acquisition, resources, supervision, writing – review & editing.

