## Supplementary Information for "KnowVolution of an efficient polyamidase through molecular dynamics simulations of incrementally docked oligomeric substrates"

### SUPPLEMENTARY MATERIALS AND METHODS

#### Supplementary materials

All chemicals used in the experiments were of analytical grade or higher quality, sourced from Merck KGaA (Darmstadt, Germany), AppliChem GmbH (Darmstadt, Germany), and Carl Roth GmbH & Co. KG (Karlsruhe, Germany). The 6-AHA dimer was obtained from Toronto Research Chemicals Inc. (North York, Canada). Enzymes and reaction buffers were supplied by New England Biolabs (Frankfurt, Germany). Oligonucleotides were acquired from Eurofins MWG Operon Inc. (Ebersberg, Germany). Plasmid extraction and PCR purification were carried out using the NucleoSpin™ Plasmid Extraction and NucleoSpin™ Gel and PCR Clean-up Kits from Macherey-Nagel GmbH & Co. KG (Düren, Germany).

#### Nucleotide and amino acid sequences of applied enzymes

Introduced mutations are underlined and in bold.

**NylC$_{p2}$-TS nucleotide sequence:**

ATGATGCATCATCATCATCATCACGGGGCAGGTGCCAATACCACACCGGTTCATGCACT
GACCGATATTGATGGTGGTATTGCAGTTGATCCGGCACCGCGTCTGGCAGGTCCGCCT
GTTTTTGGTGGTCCGGGTAATGCTGCATTCGATCTGGCACCGGTTCGTAGCACCGGTC
GTGAAATGCTGCGTTTTGATTTTCCGGGTGTTAGCATTGGTGCAGCACATTATGAAGAA
GGTCCGACAGGCGCAACCGTTATTCATATTCCGGCAGGCGCACGTACCGCAGTTGATG
CACGTGGTGGTGCAGTTGGTCTGAGCGGTGGTTATGATTTTAATCATGCAATTTGCCTG
GCAGGCGGTGCAGGTTATGGTCTGGAAGCCGGTGCCGGTGTTAGTGGTGCACTGCTG
GAACGTCTGGAATATCGTACCGGTTTTGCAGAACTGCAGCTGGTTAGCAGCGCAGTTAT
CTATGATTTTTCAGCACGTTCAACCGCAGTTTATCCTGATAAAGCACTGGGTCGTGCAG
CACTGGAATTTGCAGTTCCGGGTGAATTTCCGCAGGGTCGTGCCGGTGCGGGTATGAG
CGCAAGCGCAGGTAAAGTTGATTGGGATCGTACCGAAATTACCGGTCAGGGTGCAGCC
TTTCGTCGTCTGGGTGATGTTCGTATTCTGGCAGTTGTTGTTCCGAATCCGGTTGGTGT
TATTGTTGATCGTGCAGGCACCGTTGTTCGTGGTAATTATGATGCACAGACCGGTGTTC
GTCGTCATCCGGTTTTTGATTATCAAGAAGCATTTGCCGAACAGGTTCCTCCGGTTACC
CAAGCAGGTAATACCACAATTAGCGCCATTGTTACCAATGTGCGTATGAGTCCGGTTGA
ACTGAATCAGTTTGCGAAACAGGTTCATAGCAGCATGCATCGTGGCATTCAGCCGTTTC
ATACAGATATGGATGGTGATACCCTGTTTGCAGTTACCACCGATGAAATTGATCTGCCG
ACAACACCGGGTAGCAGCCGTGGTCGTCTGAGCGTTAATGCAACCGCACTGGGTGCAA
TTGCCAGCGAAGTTATGTGGGATGCCGTTCTGGAAGCGGGTAAATAA

**NylC$_{p2}$-TS amino acid sequence:**

MMHHHHHHGA[1]GANTTPVHALTDIDGGIAVDPAPRLAGPPVFGGPGNAAFDLAPVRSTGR
EMLRFDFPGVSIGAAHYEEGPTGATVIHIPAGARTAVDARGGAVGLSGGYDFNHAICLAGG

AGYGLEAGAGVSGALLERLEYRTGFAELQLVSSAVIYDFSARSTAVYPDKALGRAALEFAVP
GEFPQGRAGAGMSASAGKVDWDRTEITGQGAAFRRLGDVRILAVVVPNPVGVIVDRAGTV
VRGNYDAQTGVRRHPVFDYQEAFAEQVPPVTQAGNTTISAIVTNVRMSPVELNQFAKQVH
SSMHRGIQPFHTDMDGDTLFAVTTDEIDLPTTPGSSRGRLSVNATALGAIASEVMWDAVLE
AGK*

**NylC-HP nucleotide sequence:**

ATGCATCATCATCATCATCACGGGGCAGGTGCCAATACCACACCGGTTCATGCACTGA
CCGATATTGATGGTGGTATTGCAGTTGATCCGGCACCGCGTCTGGCAGGTCCGCCTGT
TTTTGGTGGTCCGGGTAATGCTGCATTCGATCTGGCACCGGTTCGTAGCACCGGTCGT
GAAATGCTGCGTTTTGATTTTCCGGGTGTTAGCATTGGTGCAGCACATTATGAAGAAGG
TCCGACAGGCGCAACCGTTATTCATATTCCGGCAGGCGCACGTACCGCAGTTGATGCA
CGTGGTGGTGCAGTTGGTCTGAGCGGTGGTTATGATTTTAATCATGCAATTTGCCTGGC
AGGCGGTGCAGGTTATGGTCTGGAAGCCGGTGCCGGTGTTAGTGGTGCACTGCTGGA
ACGTCTGGAATATCGTACCGGT**TGG**GCAGAACTGCAGCTGGTTAGCAGCGCAGTTATC
TATGATTTTTCAGCACGTTCAACCGCAGTTTATCCTGATAAAGCACTGGGTCGTGCAGC
ACTGGAATTTGCAGTTCCGGGTGAATTTCCGCAGGGTCGTGCCGGTGCGGGTATGAGC
GCAAGCGCAGGTAAAGTTGATTGGGATCGTACCGAAATTACCGGTCAGGGTGCAGCCT
TTCGTCGTCTGGGTGATGTTCGTATTCTGGCAGTTGTTGTTCCGAATCCGGTTGGTGTT
ATTGTTGATCGTGCAGGCACCGTTGTTCGTGGTAATTATGATGCACAGACCGGTGTTCG
TCGTCATCCGGTTTTTGATTATCAAGAAGCATTTGCCGAACAGGTTCCTCCGGTTACCC
AAGCAGGTAATACCACAATTAGCGCCATTGTTACCAATGTGCGTATGAGTCCGGTTGAA
CTGAATCAGTTTGCGAAACAGGTTCATAGCAGCATGCATCGTGGCATTCAGCCGTTTCA
TACA**ATG**ATGGATGGTGATACCCTGTTTGCAGTTACCACCGATGAAATTGATCTGCCGA
CAACACCGGGTAGCAGCCGTGGT**GCG**CTGAGCGTTAATGCAACCGCACTGGGTGCAA
TTGCCAGCGAAGTTATGTGGGATGCCGTTCTGGAAGCGGGTAAA

**NylC-HP amino acid sequence:**

MMHHHHHHGA[1]GANTTPVHALTDIDGGIAVDPAPRLAGQPVFGGPGNAAFDLAPVRSTGR
EMLRFDFPGVSIGAAHYEEGPTGATVIHIPAGARTAVDARGGAVGLSGGYDFNHAICLAGG
AGYGLEAGAGVSGALLERLEYRTG**W**AELQLVSSAVIYDFSARSTAVYPDKALGRAALEFAV
PGEFPQGRAGAGMSASAGKVDWDRTEITGQGAAFRRLGDVRILAVVVPNPVGVIVDRAGT
VVRGNYDAQTGVRRHPVFDYQEAFAEQVPPVTQAGNTTISAIVTNVRMSPVELNQFAKQV
HSSMHRGIQPFHT**M**MDGDTLFAVTTDEIDLPTTPGSSRG**A**LSVNATALGAIASEVMWDAVL
EAGK*

### Supplementary methods

#### Preparative scale enzyme expression and purification

Selected NylC$_{p2}$-TS variants were expressed in *E. coli* BL21 (DE3). Induction was carried out with 0.1 mM IPTG at an OD600 of 0.6, followed by a 20-hour expression at 18 °C. Cells were collected by centrifugation (3,220 x g, 20 min, 4 °C). The resulting pellet was resuspended in lysis buffer (50 mM $NaH_2PO_4$, pH 8.0, 300 mM NaCl) containing 1.5 mg/mL lysozyme and incubated for 30 min at 37 °C. Cell disruption was achieved through sonication (Vibra-Cell™ VCX 130, Sonics & Materials Inc., USA, 5 x 30 s bursts with 30 s cooling intervals, 60 % amplitude). The lysate was centrifuged (Eppendorf 5810R, 10,000 RPM, 30 min, 4 °C), filtered through a 0.45 µm syringe filter, and loaded onto a Ni-IDA 2000 column (Macherey-Nagel GmbH & Co. KG, Düren, Germany). Unspecifically bound proteins were washed away with 30 mL lysis buffer, and the target protein was eluted with 2.5 mL elution buffer (50 mM $NaH_2PO_4$, pH 8.0, 300 mM NaCl, 250 mM imidazole). The buffer was exchanged for storage buffer (50 mM Bicine, pH 8.0, 100 mM NaCl) using PD-10 columns (Cytiva, MA, USA). The protein was then concentrated using ultra-centrifugal filter units (10 kDa Amicon®, Merck KGaA, Darmstadt, Germany) and stored at −20 °C after snap freezing in liquid nitrogen. Protein purity was verified by sodium dodecyl sulfate-polyacrylamide gel electrophoresis (SDS-PAGE), and protein concentrations were measured using the Bradford method (Pierce™ Coomassie Plus, ThermoFisher Scientific, Wesel, Germany).[1]

#### Melting temperature analysis

Enzyme melting temperatures ($T_m$) of NylC$_{p2}$-TS variants were measured using nano-Differential Scanning Fluorimetry (nanoDSF) using a Prometheus NT.48 device from NanoTemper Technologies GmbH (München, Germany). Measurements were conducted at a fluorescence intensity of 45 %, with the temperature range set from 20 °C to 95 °C and a heating rate of 1 °C per minute.

#### PA crystallinity analysis by differential scanning calorimetry (DSC)

The percentage crystallinity of the PA 6 film and PA 6,6 granules was determined by DSC without and after heat treatment at different temperatures ranging from 40 to 90 °C in 10 °C increments. Prior to DSC measurement, all tested samples were dried under a vacuum overnight to remove absorbed water. Then, samples were weighed (approx. 7.4 mg for PA 6 and 10–12 mg for PA 6,6) and sealed in 50 µL aluminum pans (Perkin Elmer, part No. BO143017) with covers (Perkin Elmer, part No. BO143003). Covers were punctured (3 punctures ~0.2 mm in diameter), and encapsulated pans were weighed before and after DSC runs. DSC measurements were performed on Perkin Elmer Differential Scanning Calorimeter

DSC 8500. The instrument was calibrated for temperature and heat flow using indium standards prior to the analysis to ensure accuracy. Samples were equilibrated at 0 °C, then heated to 270 °C for PA 6 and to 310 °C for PA 6,6 at a rate of 10 °C min$^{-1}$, and cooled down by using a −50 °C cooling system, Perkin Elmer controlled Liquid Nitrogen Accessory CLN2. The area under the endothermic peak corresponding to the melting of PA 6 and PA 6,6 was integrated to determine the heat of fusion ($\Delta H_\mathrm{m}$) of the sample using Pyris Software, Version 13.3.1.0014. The crystallinity percentage was calculated using equation (S1).

$$\%\ \mathrm{Crystallinity} = \left(\frac{\Delta H_\mathrm{m}}{\Delta H_\mathrm{m}^0}\right) \times 100\% \qquad \text{Eq. (S1)}$$

Where $\Delta H_\mathrm{m}^0$ is the heat of fusion for 100% crystalline PA 6 (230 J g$^{-1}$), and PA 6,6 (190 J g$^{-1}$).[2]

**PA molecular weight analysis by gel permeation chromatography (GPC)**

Molecular weights (the number-average molecular weight ($M_\mathrm{n}$) and the weight-average molecular weight ($M_\mathrm{w}$)) and dispersity ($Đ_\mathrm{M}$) were determined by GPC. The GPC analyses were carried out using 1,1,1,3,3,3-hexafluoro-2-propanol (HFIP) (99.9 %, Chempur) as eluent, supplemented with 0.05 mol L$^{-1}$ sodium trifluoracetate (NaTFAc) (≥ 98 %, Thermo Scientific). Each investigated polymer sample was dissolved in HFIP at a concentration of 1 mg·mL$^{-1}$.

The analytical setup is equipped with an HPLC pump (1200, Agilent), a refractive index detector (RI) (1200, Agilent), and a UV-detector (VWD, 1200, Agilent). The samples contained 250 mg L$^{-1}$ 3,5-di-*tert*-4-butylhydroxytoluene (BHT, ≥99 %, Fluka) as internal standard. One pre-column (4.6 × 50 mm) and two PFG gel columns (4.6 × 250 mm, Polymer Standards Service) were applied at a flow rate of 0.3 mL min$^{-1}$ at 40 °C. The gel particles had a diameter of 5 µm, with nominal pore widths of $10^2$ and $10^3$ Å. Calibration was performed using narrowly distributed poly(methyl methacrylate) standards (Polymer Standards Service). The data analysis was conducted using the PSS WinGPC UniChrom software (Version 8.3.2).

### SUPPLEMENTARY FIGURES AND TABLES

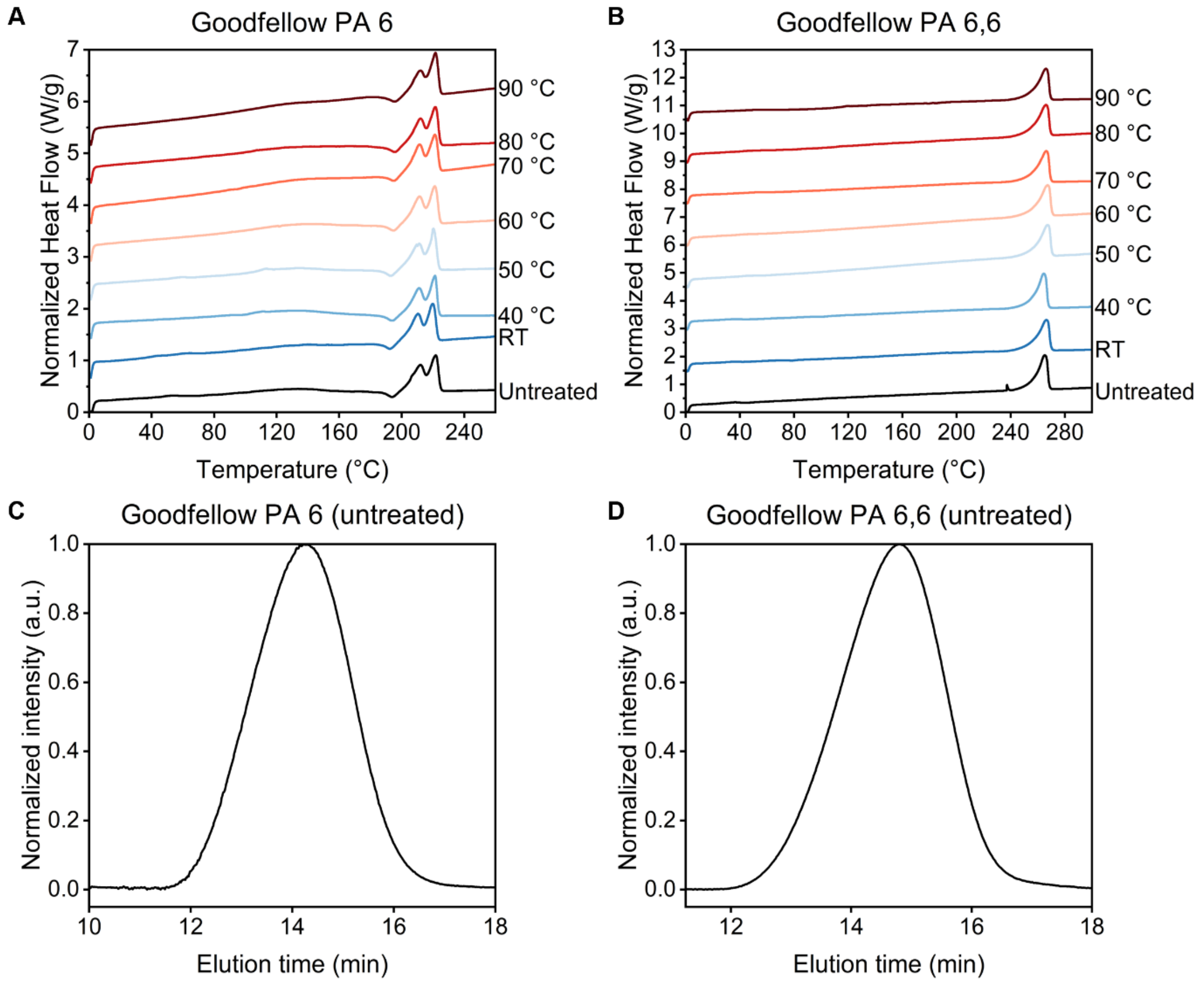

**Figure S1.** Crystallinity and molecular weight analyses of PA 6 and PA 6,6.

A) DSC thermograms of untreated and heat-treated PA 6 samples. PA 6 samples were either analyzed as provided (untreated), or pre-incubated in buffer (50 mM bicine, pH 8.0, 100 mM NaCl) at 30–90 °C for 24 h. B) Normalized GPC elugram (HFIP) of untreated Goodfellow PA 6 film (0.2 mm thick). C) Normalized GPC elugram (HFIP) of untreated Goodfellow PA 6,6 granules (3 mm diameter).

**Table S1.** Percentage crystallinity ($X_c$) of untreated and heat-treated PA 6 and PA 6,6 determined by DSC.

| **PA 6** | | | **PA 6,6** | | |
|---|---|---|---|---|---|
| Sample | $\Delta H_m$ (J g$^{-1}$) | % Crystallinity | Sample | $\Delta H_m$ (J g$^{-1}$) | % Crystallinity |
| Untreated | 49 | 21 | Untreated | 75 | 39 |
| RT | 50 | 22 | RT | 76 | 40 |
| 40 °C | 50 | 22 | 40 °C | 74 | 39 |
| 50 °C | 51 | 22 | 50 °C | 74 | 39 |
| 60 °C | 50 | 22 | 60 °C | 74 | 39 |
| 70 °C | 50 | 22 | 70 °C | 74 | 39 |
| 80 °C | 50 | 22 | 80 °C | 74 | 39 |
| 90 °C | 50 | 22 | 90 °C | 74 | 39 |

**Table S2.** Molecular weight characteristics of commercial polymer samples as determined by GPC.

This table presents the molecular weight characteristics, including number-average molecular weight ($M_n$), weight-average molecular weight ($M_w$), and dispersity index ($Đ_M$), of various commercial polymer samples.

| **Sample** | $M_n$ **(Da)** | $M_w$ **(Da)** | $Đ_M$ |
|---|---|---|---|
| Good fellow PA 6 film, 0.2 mm thickness | 44520 | 108500 | 2.438 |
| Good fellow PA 6,6 granules, 3°mm diameter | 29300 | 61840 | 2.111 |

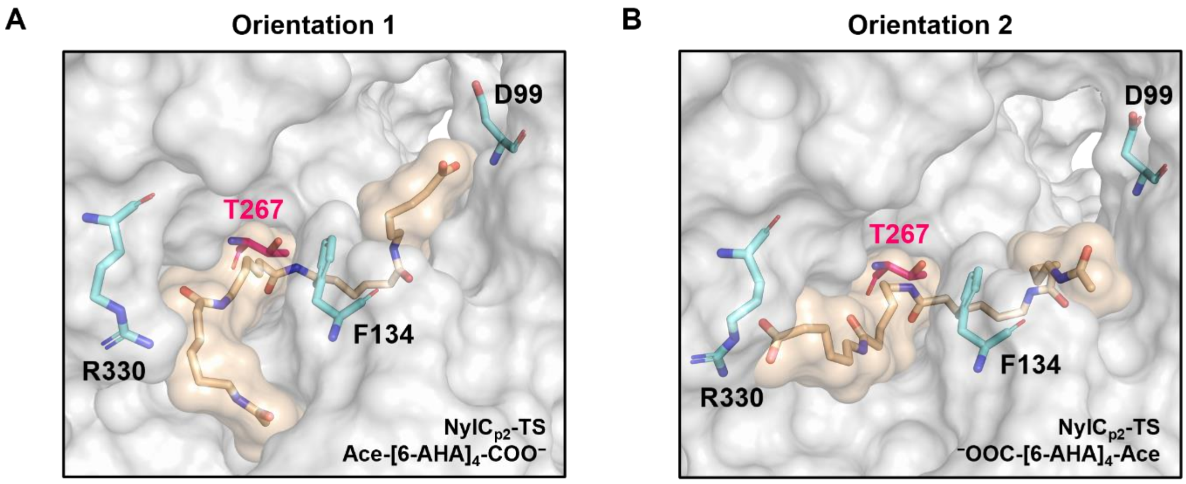

**Figure S2**. Visualization of the enzyme-substrate complexes.

A) NylC$_{p2}$-TS/Ace-[6-AHA]$_4$-COO$^-$ and B) NylC$_{p2}$-TS/$^-$OOC-[6-AHA]$_4$-Ace. As shown, the sequence of the termini (i.e., Ace and COO$^-$) used for substrate naming refers to the orientation of the substrate inside the pocket. The residues D99, F134, T267, and R330 are highlighted as reference points. The naming of the substrates NMe-[6-AHA]$_4$-$NH_3^+$ and $H_3N^+$-[6-AHA]$_4$-NMe follows the same logic.

**Table S3.** Ligand poses ranked based on their affinity score towards NylC$_{p2}$-TS.

Calculations were performed with AutoDock Vina (version 1.2.5).[3] Incremental docking of PA 6 model substrates starting from Ace-[6-AHA]-NMe and NMe-[6-AHA]-Ace revealed that the substrates Ace-[6-AHA]$_4$-COO$^-$ and $^-$OOC-[6-AHA]$_4$-Ace generate the most favorable poses (i.e., poses with low affinity scores and high populations). In this context, 6-AHA refers to 6-aminohexanoic acid, Ace to an acetyl cap, and NMe to an N-Methyl cap.

| | **Ace-[6-AHA]$_4$-COO$^-$** | | **$^-$OOC-[6-AHA]$_4$-Ace** | |
|---|---|---|---|---|
| **Rank** | Affinity (kcal mol$^{-1}$) | Occurrences out of 1000 | Affinity (kcal mol$^{-1}$) | Occurrences out of 1000 |
| 1 | −13.27 ± 0.01 | 63 | −13.81 ± 0.03 | 23 |
| 2 | −13.25 ± 0.01 | 82 | −13.71 ± 0.04 | 20 |
| 3 | −13.23 ± 0.01 | 55 | −13.59 ± 0.02 | 29 |
| 4 | −13.22 ± 0.01 | 82 | −13.58 ± 0.02 | 29 |
| 5 | −13.21 ± 0.01 | 71 | −13.55 ± 0.03 | 21 |
| 6 | −13.16 ± 0.02 | 24 | −13.48 ± 0.02 | 27 |
| 7 | −13.15 ± 0.01 | 78 | −13.47 ± 0.02 | 23 |
| 8 | −13.13 ± 0.02 | 27 | −13.42 ± 0.02 | 20 |
| 9 | −13.09 ± 0.01 | 59 | −13.36 ± 0.01 | 23 |
| 10 | −13.06 ± 0.01 | 38 | −13.33 ± 0.01 | 21 |

**Table S4**. Sequencing occurrence and specific activities of double substitutions from the iterative site-saturation mutagenesis of D304 with NylC$_{p2}$-TS$^{F134W}$ as parental gene compared to NylC$_{p2}$-TS.

Errors represent the standard error of mean.

| **Substitution F134W/D304X** | **Occurrence** | **Specific activity ($\mu mol_{6\text{-}AHAeq.}$ $h^{-1}$ $mg_{enzyme}^{-1}$)** | **Improvement (fold)** |
|---|---|---|---|
| M | 6 (13 %) | 410 ± 10 | 5.3 ± 0.1 |
| S | 6 (13 %) | 223 ± 17 | 2.9 ± 0.2 |
| E | 5 (11 %) | 239 ± 7 | 3.1 ± 0.1 |
| I | 5 (11 %) | 341 ± 5 | 4.4 ± 0.1 |
| L | 5 (11 %) | 371 ± 17 | 4.8 ± 0.2 |
| R | 5 (11 %) | 248 ± 12 | 3.2 ± 0.2 |
| V | 4 (9 %) | 202 ± 10 | 2.6 ± 0.1 |
| C | 3 (7 %) | n.d. | n.d. |
| A | 2 (4 %) | n.d. | n.d. |
| Q | 1 (2 %) | n.d. | n.d. |
| F | 1 (2 %) | n.d. | n.d. |
| T | 1 (2 %) | n.d. | n.d. |
| D (NylC$_{p2}$-TS$^{F134W}$) | 1 (2 %) | 217 ± 5 | 2.8 ± 0.1 |

**Table S5**. Conventional Michaelis-Menten kinetics of NylC$_{p2}$-TS variants for PA 6.

| **Variant** | | | | | **Conc. (nM)** | $V_{max}$ **(μM s$^{-1}$)** | $k_{cat}$ **(s$^{-1}$)** | **Specific activity (μmol$_{6\text{-AHAeq.}}$ h$^{-1}$ mg$_{enzyme}^{-1}$)** | **Improv. (fold)** |
|---|---|---|---|---|---|---|---|---|---|
| **NylC$_{p2}$-TS** | | | | | **50** | **0.040 ± 0.002** | **0.79 ± 0.03** | **75 ± 3** | **1.0** |
| | D99R | | | | 50 | 0.114 ± 0.005 | 2.27 ± 0.11 | 216 ± 10 | 2.9 |
| | | F134W | | | 50 | 0.118 ± 0.002 | 2.36 ± 0.05 | 224 ± 4 | 3.0 |
| | | | D304M | | 50 | 0.130 ± 0.001 | 2.60 ± 0.02 | 247 ± 2 | 3.3 |
| | | | | R330A | 50 | 0.077 ± 0.000 | 1.55 ± 0.01 | 147 ± 1 | 2.0 |
| | | F134W | D304M | | 25 | 0.115 ± 0.002 | 4.60 ± 0.08 | 437 ± 7 | 5.8 |
| V4 | D99R | F134W | D304M | | 25 | 0.065 ± 0.001 | 2.61 ± 0.03 | 248 ± 3 | 3.3 |
| V3 | D99G | F134W | D304M | | 25 | 0.070 ± 0.002 | 2.80 ± 0.06 | 274 ± 6 | 3.6 |
| V5 | D99V | F134W | D304M | | 25 | 0.053 ± 0.002 | 2.12 ± 0.07 | 199 ± 6 | 2.7 |
| **NylC-HP (V1)** | | **F134W** | **D304M** | **R330A** | **25** | **0.137 ± 0.005** | **5.48 ± 0.20** | **520 ± 19** | **6.9** |
| V2 | | F134W | D304M | R330Q | 25 | 0.098 ± 0.005 | 3.92 ± 0.20 | 371 ± 19 | 4.9 |
| V8 | D99R | F134W | D304M | R330A | 25 | 0.078 ± 0.001 | 3.12 ± 0.05 | 296 ± 5 | 3.9 |
| V6 | D99G | F134W | D304M | R330A | 25 | 0.108 ± 0.006 | 4.32 ± 0.24 | 410 ± 23 | 5.5 |
| V7 | D99V | F134W | D304M | R330A | 25 | 0.103 ± 0.005 | 4.12 ± 0.20 | 391 ± 19 | 5.2 |
| V10 | D99R | F134W | D304M | R330Q | 25 | 0.067 ± 0.011 | 2.68 ± 0.44 | 255 ± 42 | 3.4 |
| V9 | D99G | F134W | D304M | R330Q | 25 | 0.068 ± 0.005 | 2.72 ± 0.20 | 258 ± 19 | 3.4 |
| V11 | D99V | F134W | D304M | R330Q | 25 | 0.054 ± 0.002 | 2.16 ± 0.08 | 205 ± 8 | 2.7 |

**Table S6**. Conventional Michaelis-Menten kinetics of NylC$_{p2}$-TS variants for PA 6,6.

| **Variant** | | | | **Conc. (nM)** | $V_{max}$ **(μM s$^{-1}$)** | $k_{cat}$ **(s$^{-1}$)** | **Specific activity (μmol$_{PA6,6\ monomer\ eq.}$ h$^{-1}$ mg$_{enzyme}^{-1}$)** | **Improv. (fold)** |
|---|---|---|---|---|---|---|---|---|
| **NylC$_{p2}$-TS** | | | | **20** | **0.064 ± 0.002** | **3.20 ± 0.1** | **304 ± 9** | **1.0** |
| **NylC-HP** | **D99R** | **F134W** | **R330A** | **7.5** | **0.081 ± 0.002** | **10.8 ± 0.3** | **1026 ± 25** | **3.4** |

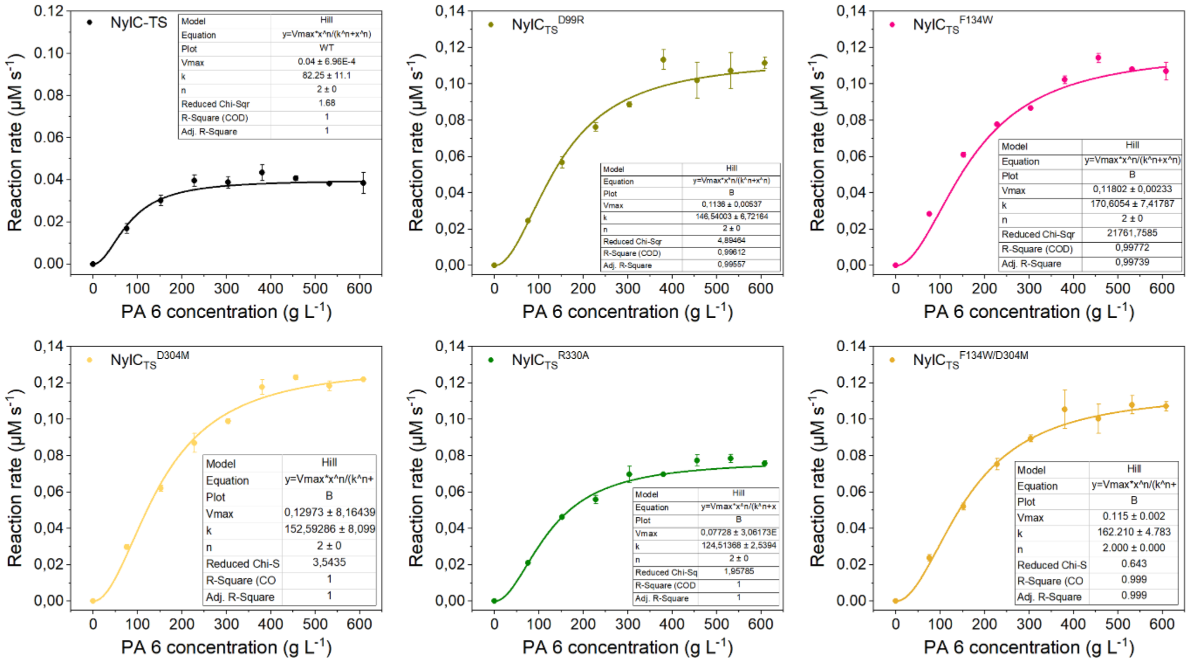

**Figure S3**. Conventional Michaelis-Menten kinetics of NylC$_{p2}$-TS, Single-substitutions and NylC$_{p2}$-TS$^{F134W/D304M}$ for PA 6.

$^{Conv}$MM kinetics were conducted at enzyme saturation (50 nM/1.90 mg L$^{-1}$ enzyme, 76–608 g L$^{-1}$ Gf-PA 6, 50 mM bicine, pH 8.0, 100 mM NaCl, 60 °C, 2 h). Reactions for individual data points were conducted in duplicate. Error bars represent the standard error of the mean.

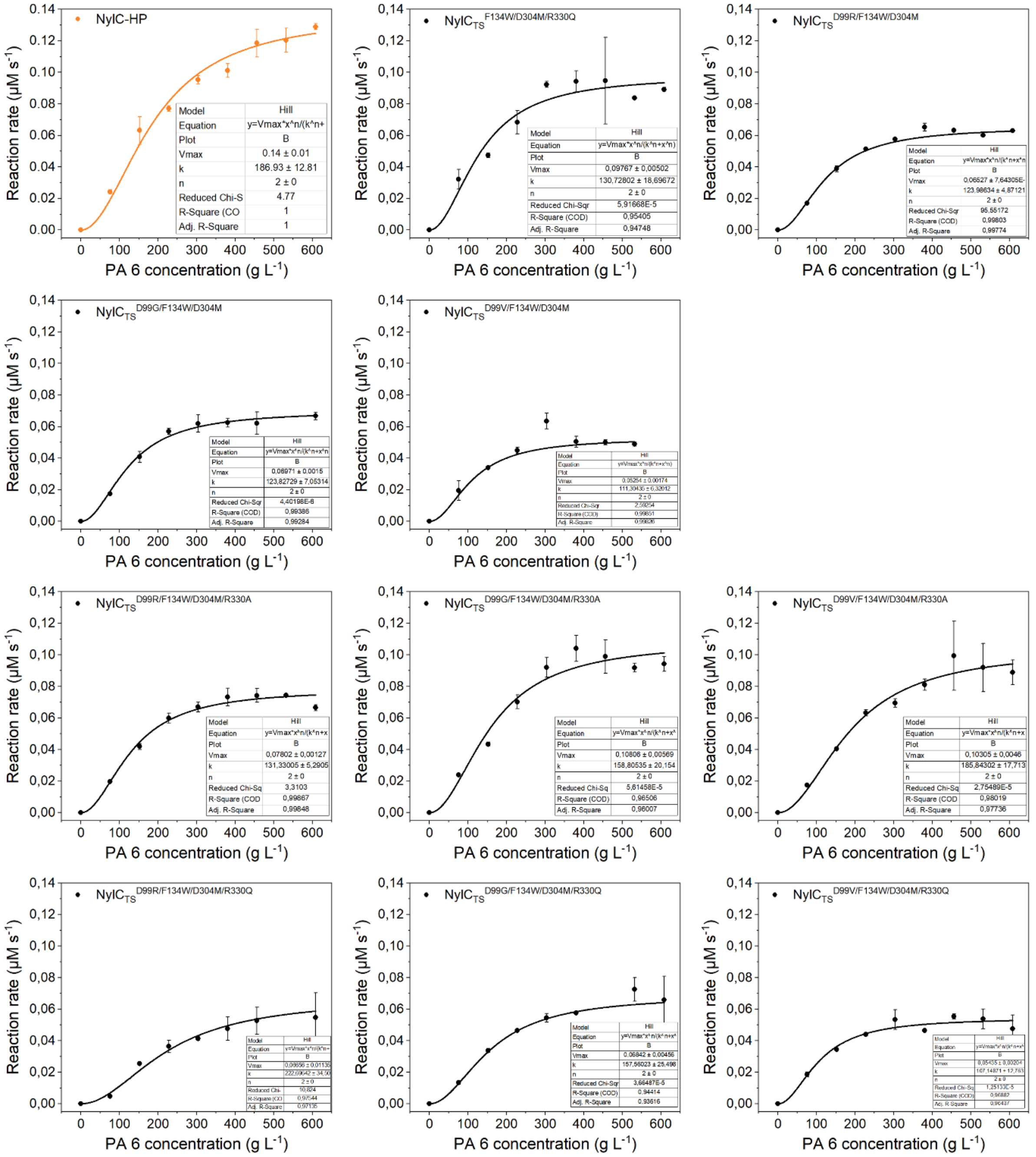

**Figure S4**. Conventional Michaelis-Menten kinetics of NylC$_{p2}$-TS triple- and quadruple-substitutions for PA 6.

$^{Conv}$MM kinetics were conducted at enzyme saturation (25 nM/0.95 mg L$^{-1}$ enzyme, 76–608 g L$^{-1}$ Gf-PA 6, 50 mM bicine, pH 8.0, 100 mM NaCl, 60 °C, 2 h). Reactions for individual data points were conducted in duplicate. Error bars represent the standard error of the mean.

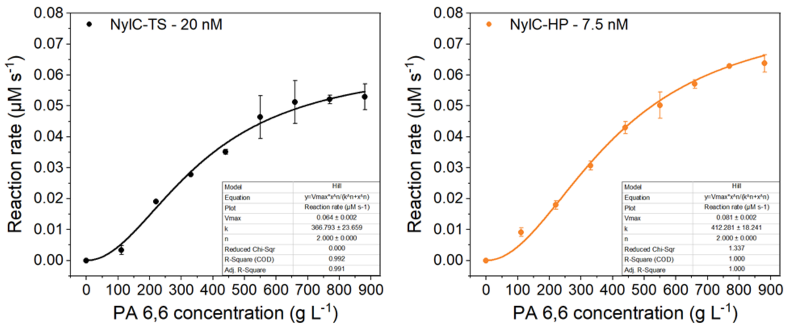

**Figure S5**. Conventional Michaelis-Menten kinetics of NylC$_{p2}$-TS and NylC-HP for PA 6,6.

$^{Conv}$MM kinetics were conducted at enzyme saturation (20 nM/0.76 mg L$^{-1}$ or 7.5 nM/0.28 mg L$^{-1}$ enzyme, 110–880 g L$^{-1}$ Gf-PA 6,6, 50 mM bicine, pH 8.0, 100 mM NaCl, 60 °C, 2 h). Reactions for individual data points were conducted in duplicate. Error bars represent the standard error of the mean.

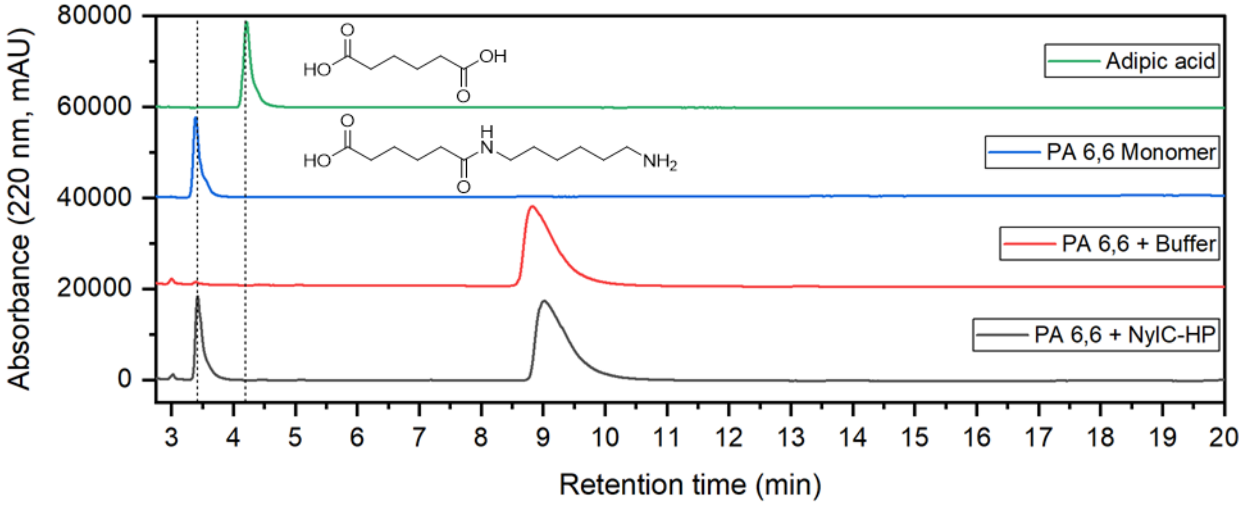

**Figure S6**. HPLC Chromatogram of adipic acid, PA 6,6 monomer, Gf-PA 6,6 control, and Gf-PA 6,6 degradation NylC-HP.

Adipic acid (green) could be resolved from PA 6,6 monomer (blue). In the control reaction with Gf-PA 6,6 but without enzyme, formation of an additional peak was observed. The degradation reaction (black; 50 nM/1.90 mg L$^{-1}$ enzyme, 660 g L$^{-1}$ Gf-PA 6, 50 mM bicine, pH 8.0, 100 mM NaCl, 60 °C, 8 h) resulted in PA 6,6 monomer as sole degradation product and no conversion of the peak extracted from Gf-PA 6,6.

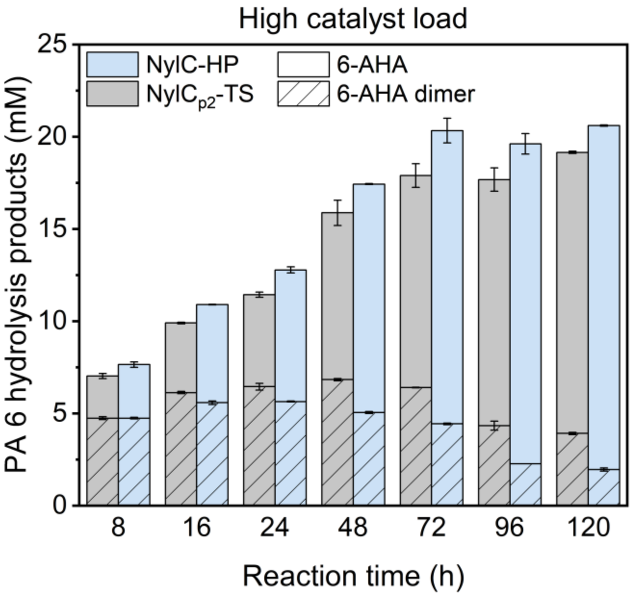

**Figure S7**. Time-resolved Gf-PA 6 degradation by NylC$_{p2}$-TS and NylC-HP with high catalyst load.

Reactions were conducted at substrate saturation (5 µM/190 mg L$^{-1}$ enzyme, 456 g L$^{-1}$ Gf-PA 6, 50 mM bicine, pH 8.0, 100 mM NaCl, 60 °C, 8–120 h).

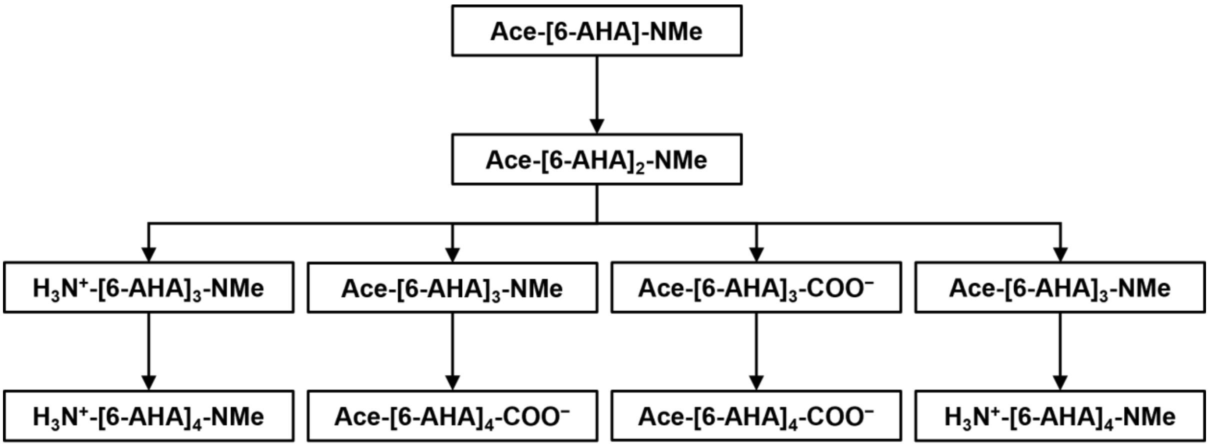

**Figure S8.** Incremental procedure for docking Ace-[6-AHA]$_4$-COO$^-$ and $H_3N^+$-[6-AHA]$_4$-NMe to the receptor.

Starting with Ace-[6-AHA]-NMe as the first fragment to be docked, both PA 6 model substrates are obtained by following each pathway characterized by a specific elongation of the fragments. In this context, 6-AHA refers to 6-aminohexanoic acid, Ace to an acetyl cap, and NMe to an N-Methyl cap.

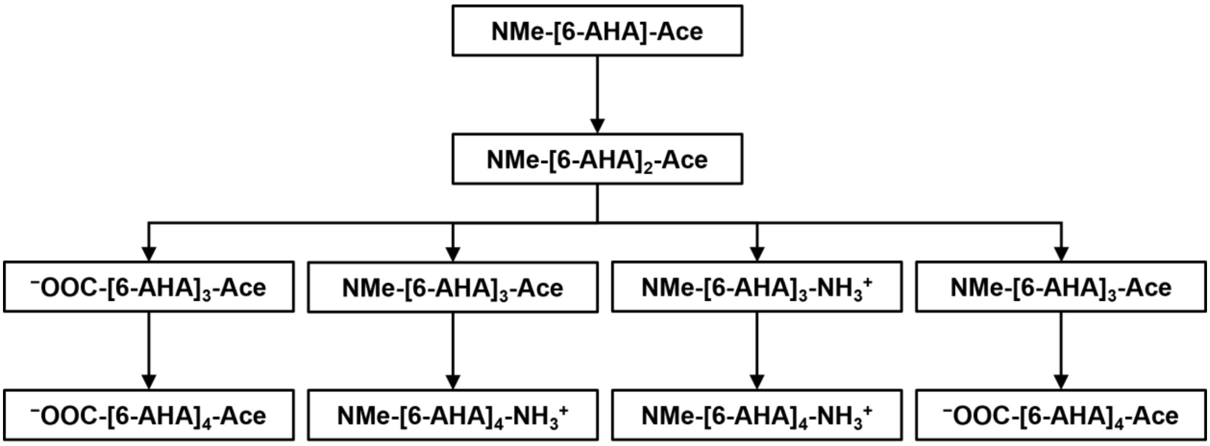

**Figure S9** Incremental procedure for docking $^-$OOC-[6-AHA]$_4$-Ace and NMe-[6-AHA]$_4$-$NH_3^+$ to the receptor.

Starting with NMe-[6-AHA]-Ace as the first fragment to be docked, both PA 6 model substrates are obtained by following each pathway characterized by a specific elongation of the fragments. In this context, 6-AHA refers to 6-aminohexanoic acid, Ace to an acetyl cap, and NMe to an N-Methyl cap.

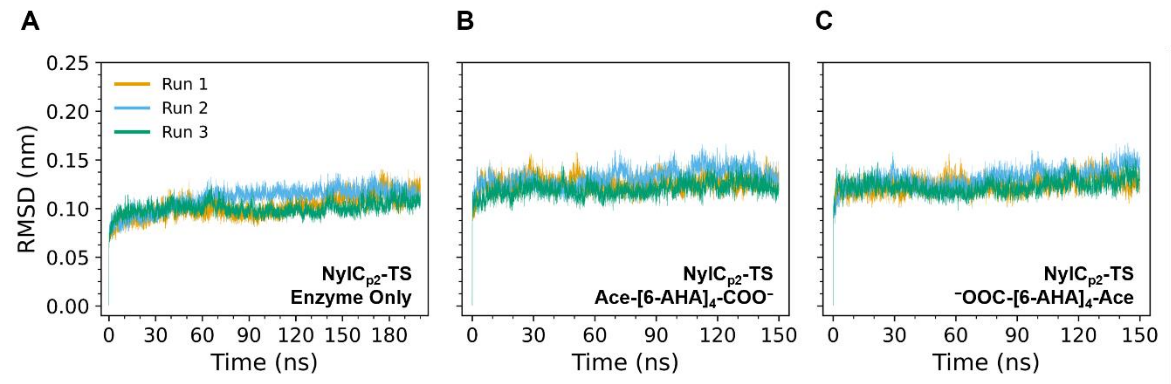

**Figure S10.** Root mean square deviation (RMSD) of the protein backbone of NylC$_{p2}$-TS.

RMSD after least squares fit to the backbone observed for molecular dynamics simulations of systems containing A) only the enzyme NylC$_{p2}$-TS or the enzyme-substrate complexes B) NylC$_{p2}$-TS/Ace-[6-AHA]$_4$-COO$^-$, and C) NylC$_{p2}$-TS/$^-$OOC-[6-AHA]$_4$-Ace.

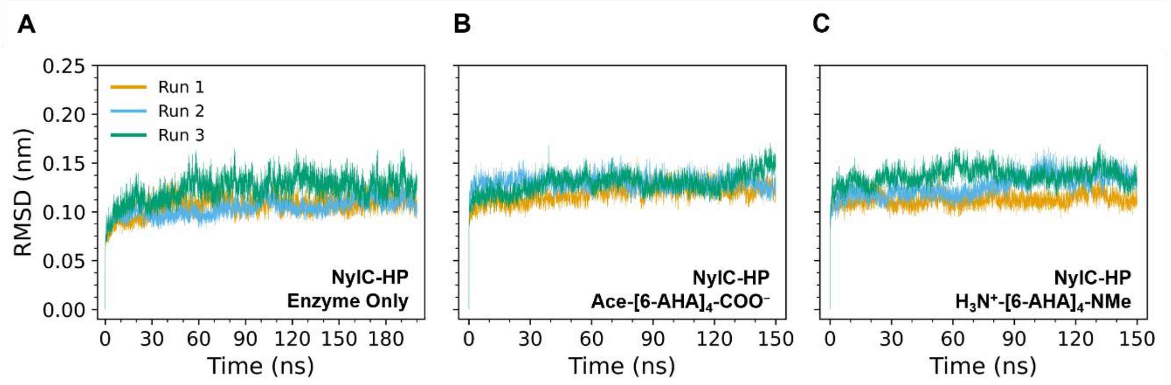

**Figure S11.** Root mean square deviation (RMSD) of the protein backbone of NylC$_{p2}$-HP.

RMSD after least squares fit to the backbone observed for molecular dynamics simulations of systems containing A) only the enzyme NylC$_{p2}$-TS$^{F134W/D304M/R330A}$ (NylC-HP) or the enzyme-substrate complexes B) NylC-HP/Ace-[6-AHA]$_4$-COO$^-$, and C) NylC-HP/$H_3N^+$-[6-AHA]$_4$-NMe.

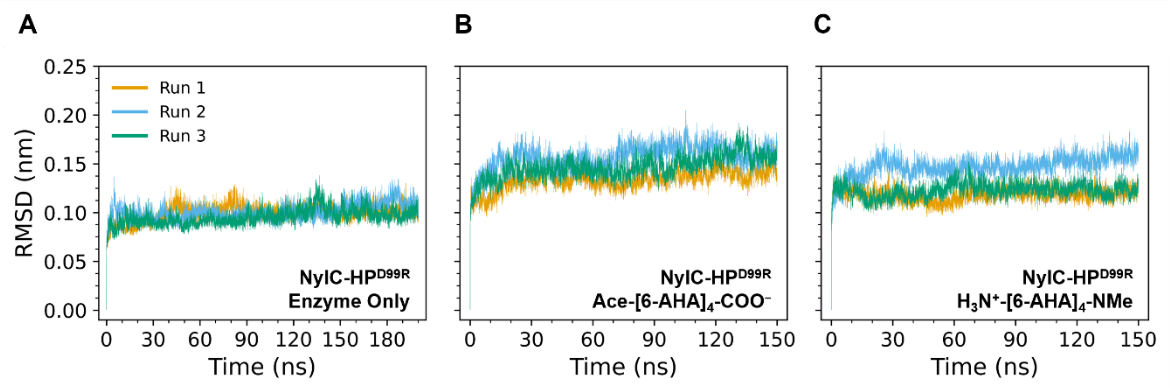

**Figure S12.** Root mean square deviation (RMSD) of the protein backbone of NylC-HP$^{D99R}$.

RMSD after least squares fit to the backbone observed for molecular dynamics simulations of systems containing A) only the enzyme NylC$_{p2}$-TS$^{D99R/F134W/D304M/R330A}$ (NylC-HP$^{D99R}$) or the enzyme-substrate complexes B) NylC-HP$^{D99R}$/Ace-[6-AHA]$_4$-COO$^-$, and C) NylC-HP$^{D99R}$/$H_3N^+$-[6-AHA]$_4$-NMe.

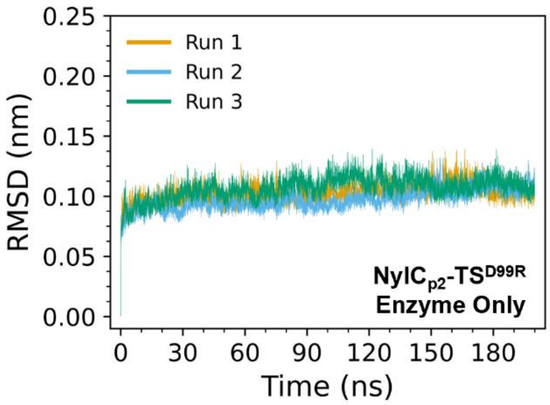

**Figure S13.** Root mean square deviation (RMSD) of the protein backbone of NylC$_{p2}$-TS$^{D99R}$.

RMSD after least squares fit to the backbone observed for molecular dynamics simulations of NylC$_{p2}$-TS$^{D99R}$.

**Table S7.** Ligand poses ranked based on their affinity score towards NylC-HP.

Calculations were performed with AutoDock Vina (version 1.2.5).[3] Incremental docking of PA 6 model substrates starting from Ace-[6-AHA]-NMe and NMe-[6-AHA]-Ace revealed that the substrates Ace-[6-AHA]$_4$-COO$^-$ and $H_3N^+$-[6-AHA]$_4$-NMe generate the most favorable poses (i.e., poses with low affinity scores and high populations). In this context, 6-AHA refers to 6-aminohexanoic acid, Ace to an acetyl cap, and NMe to an N-Methyl cap.

| | **Ace-[6-AHA]$_4$-COO$^-$** | | **$H_3N^+$-[6-AHA]$_4$-NMe** | |
|---|---|---|---|---|
| **Rank** | Affinity (kcal mol$^{-1}$) | Occurrences out of 1000 | Affinity (kcal mol$^{-1}$) | Occurrences out of 1000 |
| 1 | −13.84 ± 0.01 | 124 | −13.61 ± 0.01 | 39 |
| 2 | −13.79 ± 0.01 | 108 | −13.60 ± 0.01 | 36 |
| 3 | −13.71 ± 0.01 | 126 | −13.59 ± 0.01 | 44 |
| 4 | −13.69 ± 0.01 | 45 | −13.59 ± 0.01 | 25 |
| 5 | −13.67 ± 0.01 | 37 | −13.58 ± 0.01 | 36 |
| 6 | −13.66 ± 0.01 | 40 | −13.58 ± 0.01 | 20 |
| 7 | −13.65 ± 0.01 | 23 | −13.57 ± 0.01 | 38 |
| 8 | −13.64 ± 0.01 | 21 | −13.54 ± 0.01 | 23 |
| 9 | −13.62 ± 0.01 | 79 | −13.52 ± 0.01 | 64 |
| 10 | −13.62 ± 0.01 | 26 | −13.51 ± 0.01 | 61 |

**Table S8.** Ligand poses ranked based on their affinity score towards NylC-HP$^{D99R}$.

Calculations were performed with AutoDock Vina (version 1.2.5).[3] Incremental docking of PA 6 model substrates starting from Ace-[6-AHA]-NMe and NMe-[6-AHA]-Ace revealed that the substrates Ace-[6-AHA]$_4$-COO$^−$ and $H_3N^+$-[6-AHA]$_4$-NMe generate the most favorable poses (i.e., poses with low affinity scores and high populations). In this context, 6-AHA refers to 6-aminohexanoic acid, Ace to an acetyl cap, and NMe to an N-Methyl cap.

| | **Ace-[6-AHA]$_4$-COO$^−$** | | **$H_3N^+$-[6-AHA]$_4$-NMe** | |
|---|---|---|---|---|
| **Rank** | Affinity (kcal mol$^{−1}$) | Occurrences out of 1000 | Affinity (kcal mol$^{−1}$) | Occurrences out of 1000 |
| 1 | −13.60 ± 0.01 | 162 | −13.73 ± 0.01 | 28 |
| 2 | −13.57 ± 0.01 | 140 | −13.72 ± 0.01 | 29 |
| 3 | −13.54 ± 0.02 | 20 | −13.71 ± 0.01 | 24 |
| 4 | −13.51 ± 0.01 | 113 | −13.67 ± 0.01 | 41 |
| 5 | −13.51 ± 0.01 | 30 | −13.66 ± 0.01 | 44 |
| 6 | −13.49 ± 0.01 | 37 | −13.65 ± 0.01 | 45 |
| 7 | −13.48 ± 0.01 | 42 | −13.65 ± 0.01 | 36 |
| 8 | −13.46 ± 0.01 | 27 | −13.64 ± 0.01 | 32 |
| 9 | −13.45 ± 0.01 | 32 | −13.64 ± 0.01 | 28 |
| 10 | −13.42 ± 0.01 | 21 | −13.63 ± 0.01 | 30 |

**Table S9.** Electrostatic interactions between substrate termini and residues D99R and R330 observed during MD simulation.

Percentage of molecular dynamics (MD) simulation time with hydrogen bond ($Time_{HB}$) and salt bridge ($Time_{SB}$) existence between the carboxy-terminus of the substrate Ace-[6-AHA]$_4$-COO$^−$ and the side chain of residue R330 (chain C) of NylC$_{p2}$-TS and D99R (chain B) of NylC-HP$^{D99R}$. MD simulations were performed in triplicates for 150 ns each, with the first 20 ns not considered for analysis.

| **System** | **Residue** | **Run** | **$Time_{HB}$ (%)** | **$Time_{SB}$ (%)** |
|---|---|---|---|---|
| NylC$_{p2}$-TS / $^−$OOC-[6-AHA]$_4$-Ace | R330 | 1 | 79 | 79 |
| | | 2 | 87 | 87 |
| | | 3 | 34 | 29 |
| | | Mean | 67 ± 16 | 65 ± 18 |
| NylC-HP$^{D99R}$ / Ace-[6-AHA]$_4$-COO$^−$ | D99R | 1 | 89 | 89 |
| | | 2 | 98 | 98 |
| | | 3 | 2 | 2 |

**Table S10.** Change in interaction energy upon H-bond formation between substrate termini and residues D99R and R330 observed during MD simulation.

Mean interaction energy and standard deviation of the substrate Ace-[6-AHA]$_4$-COO$^−$ and the enzymes NylC$_{p2}$-TS and NylC-HP$^{D99R}$ with established ($E_{HB}$) and not established ($E_{no\ HB}$) hydrogen bonds (H-bonds) between the side chain of specified residues (R330 (chain C) and D99R (chain B)) of the enzymes and the carboxy-terminus of the substrate observed during molecular dynamics (MD) simulation. MD simulations were performed in triplicates for 150 ns each, with the first 20 ns not considered for analysis. As H-bond formation for MD simulation run 3 of the system NylC-HP$^{D99R}$/Ace-[6-AHA]$_4$-COO$^−$ could only be observed for 2 % of the simulation time, this value was not considered for further analysis.

| System | Residue | Run | $E_{HB}$ (kJ mol$^{−1}$) | $E_{no\ HB}$ / (kJ mol$^{−1}$) | ∆*E* / (kJ mol$^{−1}$) |
|---|---|---|---|---|---|
| NylC$_{p2}$-TS / $^−$OOC-[6-AHA]$_4$-Ace | R330 | 1 | −429 ± 40 | −337 ± 38 | −92 ± 55 |
| | | 2 | −494 ± 35 | −393 ± 27 | −101 ± 44 |
| | | 3 | −372 ± 57 | −271 ± 27 | −101 ± 63 |
| | | Mean | | | −98 ± 18 |
| NylC-HP$^{D99R}$ / Ace-[6-AHA]$_4$-COO$^−$ | D99R | 1 | −384 ± 33 | −303 ± 41 | −81 ± 53 |
| | | 2 | −421 ± 34 | −319 ± 39 | −102 ± 52 |
| | | (3 | −379 ± 45 | −377 ± 49 | −2 ± 66) |
| | | Mean | | | −92 ± 26 |

**Table S11.** Electrostatic interactions between residues D191-R330 and D99R-D304 observed during MD simulation.

Percentage of molecular dynamics (MD) simulation time with hydrogen bond (Time$_{HB}$) and salt bridge (Time$_{SB}$) existence between the side chains of specified residues (i.e., D191 (chain A) and R330 (chain C) as well as D99R (chain B) and D304 (chain A)) for the enzymes NylC$_{p2}$-TS and NylC$_{p2}$-TS$^{D99R}$. MD simulations were performed in triplicates for 200 ns each, with the first 20 ns not considered for analysis.

| System | Residues | Run | Time$_{HB}$ (%) | Time$_{SB}$ / (%) |
|---|---|---|---|---|
| NylC$_{p2}$-TS | D191-R330 | 1 | 30 | 28 |
| | | 2 | 29 | 28 |
| | | 3 | 9 | 8 |
| NylC$_{p2}$-TS$^{D99R}$ | D191-R330 | 1 | 46 | 45 |
| | | 2 | 11 | 10 |
| | | 3 | 27 | 25 |
| | D99R-D304 | 1 | 20 | 20 |
| | | 2 | 46 | 46 |
| | | 3 | 51 | 51 |

**Table S12.** Time of residue F134 being in distinct conformations for different systems containing NylC$_{p2}$-TS.

Percentage of molecular dynamics (MD) simulation time of residue F134 (chain B) being in conformation 1 (Time$_{Conf. 1}$) and conformation 2 (Time$_{Conf. 2}$) for different systems containing NylC$_{p2}$-TS. Conformation 1 is characterized by a vertically oriented side chain of F134, which prevents intramolecular interactions of the substrate. Conformation 2 is characterized by a horizontal alignment of the side chain of F134. MD simulations were performed in triplicates for 150 ns (with substrate) and 200 ns (without substrate) each, with the first 20 ns not considered for analysis.

| Substrate | Run | Time$_{Conf. 1}$ (%) | Time$_{Conf. 2}$ (%) |
|---|---|---|---|
| | 1 | 95.7 | 4.3 |
| Ace-[6-AHA]$_4$-COO$^-$ | 2 | 27.3 | 72.7 |
| | 3 | 100.0 | 0.0 |
| | 1 | 100.0 | 0.0 |
| $^-$OOC-[6-AHA]$_4$-Ace | 2 | 100.0 | 0.0 |
| | 3 | 12.0 | 88.0 |
| | 1 | 97.6 | 2.4 |
| No substrate | 2 | 96.8 | 3.2 |
| | 3 | 98.1 | 1.9 |

**Table S13.** Time of residue F134W being in distinct conformations for different systems containing NylC$_{p2}$-HP.

Percentage of molecular dynamics (MD) simulation time of residue F134W (chain B) being in conformation 1 (Time$_{Conf. 1}$) and conformation 2 (Time$_{Conf. 2}$) for different systems containing NylC$_{p2}$-TS$^{F134W/D304M/R330A}$ (NylC-HP). Conformation 1 is characterized by a vertically oriented side chain of F134W, which prevents intramolecular interactions and (optionally) fixes the amide bond to be cleaved by H-bonding close to the active residue T267. Conformation 2 is characterized by a horizontal alignment of the side chain of F134W. MD simulations were performed in triplicates for 150 ns (with substrate) and 200 ns (enzyme only) each, with the first 20 ns not considered for analysis.

| Substrate | Run | Time$_{Conf. 1}$ (%) | Time$_{Conf. 2}$ (%) |
|---|---|---|---|
| | 1 | 0.0 | 100.0 |
| Ace-[6-AHA]$_4$-COO$^-$ | 2 | 100.0 | 0.0 |
| | 3 | 34.7 | 65.3 |
| | 1 | 89.4 | 10.6 |
| $H_3N^+$-[6-AHA]$_4$-NMe | 2 | 100.0 | 0.0 |
| | 3 | 100.0 | 0.0 |
| | 1 | 99.7 | 0.3 |
| No substrate | 2 | 100.0 | 0.0 |
| | 3 | 100.0 | 0.0 |

**Table S14.** Time of residue F134W being in distinct conformations for different systems containing NylC-HP$^{D99R}$.

Percentage of molecular dynamics (MD) simulation time of residue F134W (chain B) being in conformation 1 ($Time_{Conf.\,1}$) and conformation 2 ($Time_{Conf.\,2}$) for different systems containing $NylC_{p2}$-TS$^{D99R/F134W/D304M/R330A}$ (NylC-HP$^{D99R}$). Conformation 1 is characterized by a vertically oriented side chain of F134W, which prevents intramolecular interactions and (optionally) fixes the amide bond to be cleaved by H-bonding close to the active residue T267. Conformation 2 is characterized by a horizontal alignment of the side chain of F134W. MD simulations were performed in triplicates for 150 ns (with substrate) and 200 ns (enzyme only) each, with the first 20 ns not considered for analysis.

| Substrate | Run | $Time_{Conf.\,1}$ (%) | $Time_{Conf.\,2}$ (%) |
|---|---|---|---|
| | 1 | 100.0 | 0.0 |
| Ace-[6-AHA]$_4$-COO$^-$ | 2 | 79.7 | 20.3 |
| | 3 | 100.0 | 0.0 |
| | 1 | 100.0 | 0.0 |
| $H_3N^+$-[6-AHA]$_4$-NMe | 2 | 100.0 | 0.0 |
| | 3 | 100.0 | 0.0 |
| | 1 | 100.0 | 0.0 |
| No substrate | 2 | 99.6 | 0.4 |
| | 3 | 100.0 | 0.0 |

**Table S15.** Percentage of molecular dynamics (MD) simulation time of residue F134 (chain B) being in conformation 1 ($Time_{Conf.\,1}$) and conformation 2 ($Time_{Conf.\,2}$) for different systems containing $NylC_{p2}$-TS$^{D99R}$.

Conformation 1 is characterized by a vertically oriented side chain of F134. Conformation 2 is characterized by a horizontal alignment of the side chain of F134. MD simulations were performed in triplicates for 200 ns each, with the first 20 ns not considered for analysis.

| Substrate | Run | $Time_{Conf.\,1}$ (%) | $Time_{Conf.\,2}$ (%) |
|---|---|---|---|
| | 1 | 99.4 | 0.6 |
| No substrate | 2 | 93.5 | 6.5 |
| | 3 | 35.6 | 64.4 |

**Table S16.** H-bonds between the substrate and residue at position 134 observed during MD simulation.

Percentage of molecular dynamics (MD) simulation time with hydrogen bond existence between the side chain of residue at position 134 (chain B) for different enzymes (i.e., NylC$_{p2}$-TS, NylC-HP, and NylC-HP$^{D99R}$) and the amide bond to be cleaved of various substrates (i.e., Ace-[6-AHA]$_4$-COO$^-$, $^-$OOC-[6-AHA]$_4$-Ace, and $H_3N^+$-[6-AHA]$_4$-NMe). MD simulations were performed in triplicates for 200 ns each, with the first 20 ns not considered for analysis.

| System with X = [6-AHA]$_4$ | | | | | | |
|---|---|---|---|---|---|---|
| **Enzyme** | **NylC$_{p2}$-TS** | | **NylC-HP** | | **NylC-HP$^{D99R}$** | |
| **Run** | Ace-X-COO$^-$ | $^-$OOC-X-Ace | Ace-X-COO$^-$ | $H_3N^+$-X-NMe | Ace-X-COO$^-$ | $H_3N^+$-X-NMe |
| 1 | 0 | 0 | 0 | 25 | 1 | 16 |
| 2 | 0 | 0 | 6 | 8 | 0 | 0 |
| 3 | 0 | 0 | 0 | 5 | 0 | 12 |

**Table S17.** List of primers

| Primer name | Primer sequence |
|---|---|
| pET21a_NylC$_{p2}$-TS_SSM V18_fwd | GTGGTATTGCA**NNK**GATCCGGCACCGCGTCTG |
| pET21a_NylC$_{p2}$-TS_SSM V18_rev | CATCAATATCGGTCAGTGCATGAACCGGTGTGGTATTGG |
| pET21a_NylC$_{p2}$-TS_SSM L24_fwd | CACCGCGT**NNK**GCAGGTCCGCCTGTTTTTG |
| pET21a_NylC$_{p2}$-TS_SSM L24_rev | CCGGATCAACTGCAATACCACCATCAATATCGGTCAGTG |
| pET21a_NylC$_{p2}$-TS_SSM P27_fwd | GTGGTCCGGGTAATGCTGCATTC |
| pET21a_NylC$_{p2}$-TS_SSM P27_rev | CAAAAACAGG**MNN**ACCTGCCAG |
| pET21a_NylC$_{p2}$-TS_SSM F38_fwd | CGGTTCGTAGCACCGGTCGTG |
| pET21a_NylC$_{p2}$-TS_SSM F38_rev | GTGCCAGATC**MNN**TGCAGCATTACCC |
| pET21a_NylC$_{p2}$-TS_SSM A91_fwd | CACGTGGTGGT**NNK**GTTGGTCTGAGC |
| pET21a_NylC$_{p2}$-TS_SSM A91_rev | CATCAACTGCGGTACGTGCGCCTGCC |
| pET21a_NylC$_{p2}$-TS_SSM Y98_fwd | CAGTTGGTCTGAGCGGTGGT**NNK**GATTTTAATCATGCAATTTGCC |
| pET21a_NylC$_{p2}$-TS_SSM Y98_rev | CACCACCACGTGCATCAACTGCGGTACG |
| pET21a_NylC$_{p2}$-TS_SSM D99_fwd | GTCTGAGCGGTGGTTAT**NNK**TTTAATCATGCAATTTGC |
| pET21a_NylC$_{p2}$-TS_SSM D99_rev | CAACTGCACCACCACGTGCATCAACTG |
| pET21a_NylC$_{p2}$-TS_SSM F100_fwd | GCAGGCGGTGCAGGTTATGG |
| pET21a_NylC$_{p2}$-TS_SSM F100_rev | CAGGCAAATTGCATGATT**MNN**ATCATAACCACCGCTC |
| pET21a_NylC$_{p2}$-TS_SSM G111_fwd | GCGGTGCA**NNK**TATGGTCTGGAAGCCGG |
| pET21a_NylC$_{p2}$-TS_SSM G111_rev | CTGCCAGGCAAATTGCATGATTAAAATCATAACCACCGC |
| pET21a_NylC$_{p2}$-TS_SSM F134_fwd | GAATATCGTACCGGT**NNK**GCAGAACTGCAGCTGG |
| pET21a_NylC$_{p2}$-TS_SSM F134_rev | CAGACGTTCCAGCAGTGCACCACTAACACC |
| pET21a_NylC$_{p2}$-TS_SSM L137_fwd | CGGTTTTGCAGAA**NNK**CAGCTGGTTAGCAGC |
| pET21a_NylC$_{p2}$-TS_SSM L137_rev | TACGATATTCCAGACGTTCCAGCAGTGCACCAC |
| pET21a_NylC$_{p2}$-TS_SSM L139_fwd | GTTTTGCAGAACTGCAG**NNK**GTTAGCAGCGC |
| pET21a_NylC$_{p2}$-TS_SSM L139_rev | CGGTACGATATTCCAGACGTTCCAGCAGTGCACC |
| pET21a_NylC$_{p2}$-TS_SSM V144_fwd | GTTAGCAGCGCA**NNK**ATCTATGATTTTTCAGCAC |
| pET21a_NylC$_{p2}$-TS_SSM V144_rev | CAGCTGCAGTTCTGCAAAACCGGTAC |
| pET21a_NylC$_{p2}$-TS_SSM Y146_fwd | GTTAGCAGCGCAGTTATC**NNK**GATTTTTCAGCACGTTCAAC |
| pET21a_NylC$_{p2}$-TS_SSM Y146_rev | CAGCTGCAGTTCTGCAAAACCGGTACGATATTCC |
| pET21a_NylC$_{p2}$-TS_SSM A160_fwd | GTTTATCCTGATAAA**NNK**CTGGGTCGTGCAGCACTGG |
| pET21a_NylC$_{p2}$-TS_SSM A160_rev | TGCGGTTGAACGTGCTGAAAAATCATAGATAACTGCGCTG |
| pET21a_NylC$_{p2}$-TS_SSM D209_fwd | CAGTTGTTGTTCCGAATCCGGTTGGTG |
| pET21a_NylC$_{p2}$-TS_SSM D209_rev | CCAGAATACGAAC**MNN**ACCCAGACGACG |
| pET21a_NylC$_{p2}$-TS_SSM F301_fwd | GTGATACCCTGTTTGCAGTTACCACCG |
| pET21a_NylC$_{p2}$-TS_SSM F301_rev | CATCCATATCTGTATG**MNN**CGGCTGAATGCCACG |
| pET21a_NylC$_{p2}$-TS_SSM D304_fwd | CCGTTTCATACA**NNK**ATGGATGGTGATACCCTGTTTGC |
| pET21a_NylC$_{p2}$-TS_SSM D304_rev | CTGAATGCCACGATGCATGCTGCTATGAAC |
| pET21a_NylC$_{p2}$-TS_SSM M305_fwd | CAGTTACCACCGATGAAATTGATCTGCCG |

| | |
|---|---|
| pET21a_NylC$_{p2}$-TS_SSM M305_rev | CAAACAGGGTATCACCATC**MNN**ATCTGTATGAAACGG |
| pET21a_NylC$_{p2}$-TS_SSM R330_fwd | GTAGCAGCCGTGGT**NNK**CTGAGCGTTAATGCAAC |
| pET21a_NylC$_{p2}$-TS_SSM R330_rev | CCGGTGTTGTCGGCAGATCAATTTCATCGGTGG |
| pET21a_NylC$_{p2}$-TS_SDM D99G_fwd | GGTTAT**GGT**TTTAATCATGCAAT |
| pET21a_NylC$_{p2}$-TS_SDM D99G_rev | ATTAAA**ACC**ATAACCACCGCTCAG |
| pET21a_NylC$_{p2}$-TS_SSM D99V_fwd | GGTTAT**GTT**TTTAATCATGCAAT |
| pET21a_NylC$_{p2}$-TS_SSM D99V_rev | ATTAAA**AAC**ATAACCACCGCTCAG |
| pET21a_NylC$_{p2}$-TS_SDM D99R_fwd | GCAGTTGGTCTGAGCGGTGGTTAT**CGC**TTTAATCATGCAATTTGC |
| pET21a_NylC$_{p2}$-TS_SDM D99R_rev | ACCACCACGTGCATCAACTGCGGTACGTG |
| pET21a_NylC$_{p2}$-TS_SDM F134W_fwd | GAATATCGTACCGGT**TGG**GCAGAACTGCAGCTGG |
| pET21a_NylC$_{p2}$-TS_SDM F134W_rev | CAGACGTTCCAGCAGTGCACCACTAACACC |
| pET21a_NylC$_{p2}$-TS_SDM D304M_fwd | CCGTTTCATACA**ATG**ATGGATGGTGATACCCTGTTTGC |
| pET21a_NylC$_{p2}$-TS_SDM D304_rev | CTGAATGCCACGATGCATGCTGCTATGAAC |
| pET21a_NylC$_{p2}$-TS_SSM R330A_fwd | GTAGCAGCCGTGGT**GCG**CTGAGCGTTAATGCAAC |
| pET21a_NylC$_{p2}$-TS_SDM R330_rev | CCGGTGTTGTCGGCAGATCAATTTCATCGGTGGTAAC |
| pET21a_NylC$_{p2}$-TS_SDM R330Q_fwd | CGTGGT**CAG**CTGAGCGTTAATGC |
| pET21a_NylC$_{p2}$-TS_SDM R330Q_rev | GCTCAG**CTG**ACCACGGCTGCTACC |
